# Chalcone isomerase-like impedes the lactone shunt and enhances flux partitioning in a bifurcated pathway towards isoflavonoid biosynthesis

**DOI:** 10.1101/2025.02.11.637686

**Authors:** Lee Marie Raytek, Brandon Corey Saltzman, Meha Sharma, Soon Goo Lee, Mehran Dastmalchi

## Abstract

The reconstitution of biosynthetic pathways in heterologous hosts is often challenged by the switch to a foreign cellular environment, lacking compatible structural or regulatory features. Auxiliary or non-catalytic proteins can play a critical role in modulating metabolic flux and pathway efficiency. Chalcone isomerase-like (CHIL) is a non-catalytic protein known to serve as a partner to chalcone synthase (CHS) in flavonoid biosynthesis, rectifying its promiscuous activity and preventing by-product formation, such as the aberrant *p*-coumaroyltriacetic acid lactone (CTAL). Here, we extended the characterization of CHILs to the legume-characteristic isoflavonoid pathway. We assessed four CHIL orthologs from diverse plant lineages: *Glycine max* (GmCHIL), *Oryza sativa* (OsCHIL), *Selaginella moellendorffii* (SmCHIL), and *Marchantia polymorpha* (MpCHIL). Structural modelling suggested that naringenin (flavanone) entry into the CHIL binding cleft may be sterically hindered compared to catalytic CHIs. Moreover, legume CHIL isoforms possess an additional bulky residue, Tyr^48^, that is expected to impose further constraints on ligand binding. *In vitro,* CHS produced up to 60% lactone CTAL instead of its desired output; however, CHIL suppressed this aberrant activity to 10%, concomitantly increasing target compound titers. Combinatorial enzyme and yeast biotransformation assays revealed a critical role for CHIL in conducting flux through chalcone, flavanone, and isoflavone biosynthesis. The inclusion of CHIL in our engineered yeast strains enhanced overall titers and, unexpectedly, promoted carbon flux toward the so-called deoxy-branch (isoliquiritigenin, liquiritigenin, and daidzein) by up to 67%, with a 33% increase in final daidzein titers. By extending CHIL characterization to the isoflavonoid pathway, we have revealed an expanded role for this auxiliary protein and underscored its utility in engineered metabolic contexts. Our findings reiterate the often-overlooked impact of non-catalytic proteins in shaping specialized metabolism.

**HIGHLIGHTS:**

- Combinatorial enzyme assays reveal a species-dependent preference for CHILs from more closely related plant phyla by soybean CHS, which improved chalcone and downstream flavanone output by suppressing aberrant lactone formation.
- Yeast co-expressing CHIL with the components of the isoflavonoid metabolon exhibited a 67% increase in flux through a legume-characteristic branch of the pathway, resulting in a 33% increase in titers of the major isoflavone, daidzein.
- CHIL proteins can be included to engineer the output of phenylpropanoid-derived intermediates preferentially toward isoflavonoid biosynthesis.
- Auxiliary components, such as CHIL, can be designed to refine metabolic composition in bifurcated pathways, as well as enhancing general flux through pathways.

## INTRODUCTION

Chalcone scaffolds are the entry point for thousands of downstream (iso)flavonoids through highly diverging pathways across all land plants (embryophytes). Flavonoids are plant natural products with antioxidant, anti-inflammatory, and anticancer properties and have been associated with lowering the risk or mitigating the progression of cardiovascular disease, Alzheimer’s disease, and various cancers (1–3). The structurally related isoflavonoids are characteristic of the Fabaceae (legume family), exhibiting similar broad-spectrum biological activity (4). Isoflavonoids serve critical physiological roles, including facilitating plant-microbe signalling in beneficial rhizobial symbiosis and defending plants against bacterial and fungal pathogens (5–7). Reconstructing the pathway to (iso)flavonoids in microbial chassis would enable more efficient synthesis and purification for applications, including nutraceuticals, medicines, and agrochemicals (8). Further, microbial reconstitution opens access to chemistries beyond the plant catalogue by incorporating new-to-nature derivatization (9, 10). However, achieving efficient bioproduction requires complete resolution of biosynthetic pathways and, importantly, strategies for flux enhancement to bypass bottlenecks and avoid undesirable shunts.

The committed steps of the phenylpropanoid and flavonoid pathways, phenylalanine ammonia-lyase (PAL; EC 4.3.1.24) and the type III polyketide synthase, chalcone synthase (CHS; EC 2.3.1.74), respectively, are closely linked to the migration of early plant species onto land (11). In a four-step reaction, CHS catalyzes the serial addition of three acetyl groups from malonyl-CoA substrates onto *p*-coumaroyl-CoA, followed by an intramolecular C6/C1 Claisen cyclization, forming naringenin chalcone (NC; 2′,4,4′,6′-tetrahydroxychalcone) (Fig. 1) (12). An aldo-keto reductase known as chalcone reductase (CHR; EC 2.3.1.170) appears to have been recruited much later within the Fabaceae family and acts in concert with CHS to produce a mix of NC and the 6′-deoxychalcone counterpart, isoliquiritigenin (ISO; 4,2′,4′-trihydroxychalcone) (13). The joint activity of CHS-CHR bifurcates the pathway into parallel branches, which are then acted upon by chalcone isomerase (CHI; EC 5.5.1.6). All vascular land plants possess type 1 CHIs, which perform a stereospecific isomerization of NC into (2*S*)-naringenin (NAR; 4′,5,7-trihydroxyflavanone) (14, 15). In addition, type 2 CHI activity has been noted sporadically throughout plant phyla, particularly in the Fabaceae, using both NC and ISO to produce their respective flavanones, NAR and (2*S*)-liquiritigenin (LIQ; 4′,7-dihydroxyflavanone) (14–17). Subsequently, the legume-specific cytochrome P450, isoflavone synthase (IFS; CYP93C; EC 1.14.14.87), performs a C2 to C3 aryl-ring migration to form the isoflavone skeleton (18). The resulting isoflavones, genistein (GEN; 4′,5,7-trihydroxyisoflavone) and its 5-deoxyisoflavone counterpart daidzein (DZN; 4′,7-dihydroxyisoflavone), serve as scaffolds for diverse lineage-specific tailoring. In particular, DZN is a valuable target and a precursor for potent antimicrobial pterocarpans.

**Fig. 1.**
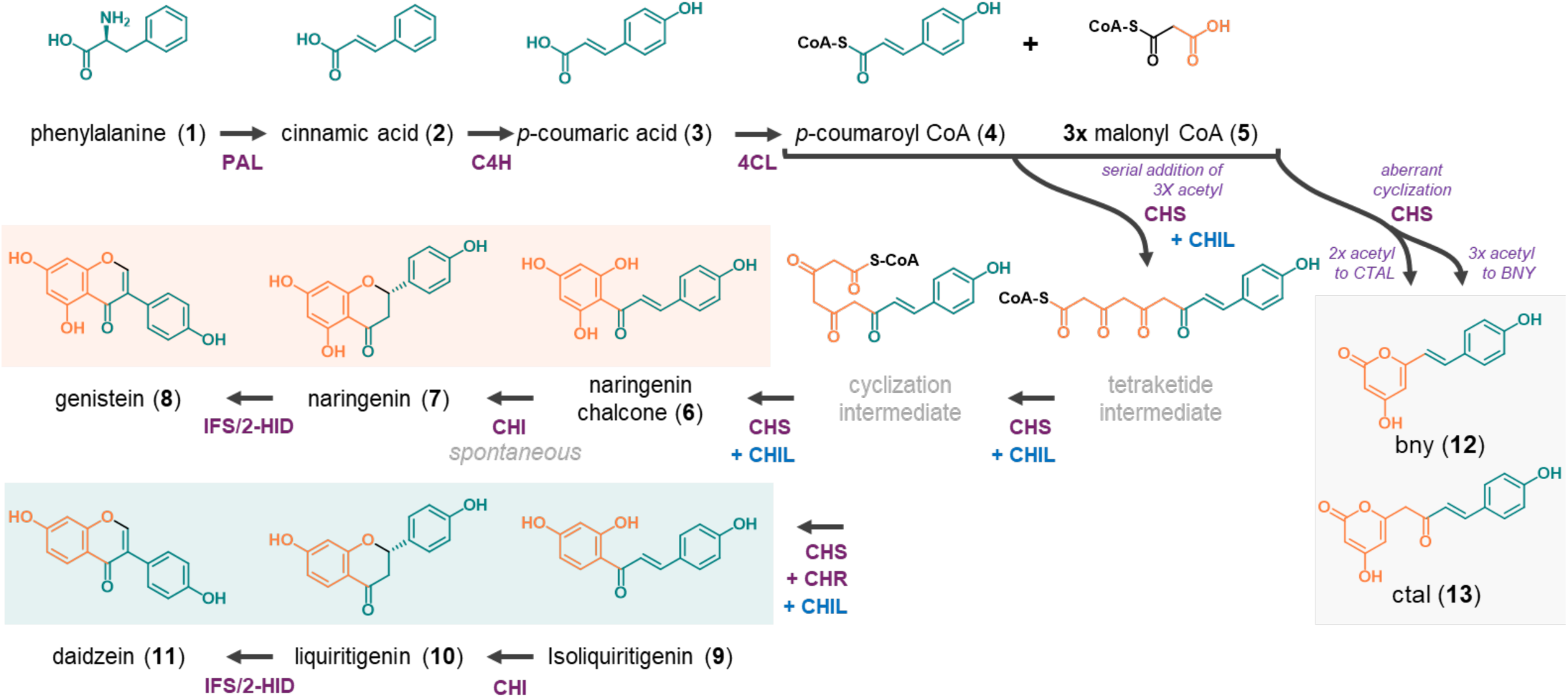
The bifurcated pathway from phenylalanine to isoflavones. Beginning with the shikimate-derived (*L*)-phenylalanine (shown in teal), the consecutive action of phenylalanine ammonia lyase (PAL), cinnamate 4-hydroxylase (C4H), and 4-coumarate-CoA ligase (4CL) yields *p*-coumaroyl-CoA. Chalcone synthase (CHS) catalyzes a four-step reaction to add three acetyl groups (shown in orange) from malonyl-CoA onto *p*-coumaroyl-CoA, generating the chalcone scaffold. In grey, the lactone shunt or by-products of CHS activity are shown, which include bis-noryangonin (BNY) and p-coumaroyltriacetic acid lactone (CTAL). The joint activity of CHS and chalcone reductase (CHR) leads to the bifurcation of the pathway and the “deoxy-branch.” Chalcone isomerase (CHI), isoflavone synthase (IFS), and 2-hydroxyisoflavanone dehydratase (2-HID) act in parallel pathways to ultimately yield genistein or daidzein. Enzymatic proteins are indicated in purple, and the auxiliary protein CHIL is shown in blue. The latter is speculated to function in tandem with CHS or the CHS-CHR complex.

A major hurdle to efficient CHS biosynthesis of chalcones is the undesired lactone shunt, which occurs due to premature cyclization of the elongating polyketide intermediates (Fig. 1). The canonical pathway is derailed after the addition of either two or three acetyl groups, yielding bis-noryangonin (BNY) or *p*-coumaroyltriacetic acid lactone (CTAL), respectively. CHS promiscuity only becomes evident *ex planta*, wherein *in vitro* or heterologous *in vivo* (e.g., *Escherichia coli, Saccharomyces cerevisiae*) assays produce these by-products at the expense of chalcones (17, 19–21). The promiscuity of ancestral plant type III PKSs, combined with the structural modifications of these homodimeric enzymes, has led to the diversification of more than twenty functionally different paralogs of CHS (e.g., stilbene synthase, benzophenone synthase, curcumin synthase) (22–24). This expansion of the enzyme family has driven the evolution of impressive diversity in the molecular scaffolds of plant polyketides (25). However, it is notable that the lactone by-products of CHS have never been reported in the plant.

Initially identified as enhancers of flavonoid production (EFP), chalcone isomerase-like (CHIL) proteins, or type 4 CHIs, function as non-catalytic, auxiliary partners to CHS, repressing the lactone shunt and increasing NC production *in vitro* and *in planta* (15, 21, 26, 27). Biochemical characterization has revealed that CHIL rectifies CHS promiscuity without increasing overall enzyme activity, whereas catalytically active CHI isoforms (types 1 and 2) do not affect by-product formation. Intriguingly, evolution of the plant CHI-fold protein superfamily appears to have followed a trajectory from fatty acid binding proteins (FAPs; type 3), with large non-polar cavities sequestering the aliphatic chains of fatty acids, to CHILs (type 4) and CHIs (types 1 and 2), which feature sterically-restricted pockets lined with polar residues to form hydrogen-bond networks (28–31). In CHIs, this hydrogen-bond network is critical for binding and stabilizing chalcones/flavanones in the active site. This neofunctionalization of a catalytic enzyme from a non-catalytic ancestor is a striking example of molecular evolution, where structural adaptations in the active site enabled a shift from passive binding to active catalysis (29). Moreover, CHIL binds CHS with strong affinity (*K*_D_ 1-126 nM) and also interacts with cytochrome P450s involved in (iso)flavonoid pathways (i.e. IFS and flavone synthase II), which act as hubs for biosynthetic complexes on the surface of the endoplasmic reticulum (ER) (21, 32). While CHIL has not been probed for binary interactions with other core (iso)flavonoid enzymes, such as CHI and CHR, it may play a role within a larger metabolon.

Non-catalytic type 4 CHILs play a crucial role in regulating CHS-catalyzed reactions, but their influence on the legume-specific 6′-deoxychalcone branch and isoflavonoid biosynthesis has remained unexplored. Through multi-enzyme assays, we demonstrated that CHIL inhibited aberrant lactonization of CHS intermediates, reducing CTAL levels from 60% to 10% and consequently increasing NAR production by up to 2-fold. In yeast, incorporating GmCHIL boosted total product accumulation by 75%, with the final isoflavone concentration reaching 14 μM/OD from 250 μM p-coumaric acid. Furthermore, DZN concentrations increased by at least 33% in all CHIL-expressing strains. Selecting superior isoforms significantly impacted flux, with optimized 4CL increasing pathway entry 4-fold, while GmCHIL and OsCHIL enhanced flavanone and isoflavone production. Surprisingly, CHIL proteins directed carbon flux preferentially into the deoxy-branch, amplified by the addition of IFS. Overall, CHIL appears to play a key role in organizing and partitioning branch specificity. This finding highlights the importance of auxiliary or structural elements in biosynthetic modules, which can enhance flux and integrate seamlessly with other metabolic engineering strategies.

## RESULTS

### Comparative sequence analysis of CHI-fold proteins in plants

Chalcone isomerase-fold amino acid sequences were compiled from diverse plant phyla by querying each sub-type using the corresponding soybean CHIs against the Phytozome database. Subsequently, alignments were generated comparing 262 type 1, 55 type 2, and 220 type 4 CHI-fold orthologs at residues known to be significant for type 1 or 2 catalytic activity or ligand-binding (Fig. 2A and Supplementary Fig. 1). Basal bryophyte/lycophyte type 2 CHI sequences were not considered due to the comparatively low number of verified, active enzymes in this clade. Our global amino acid analysis revealed that, in contrast to the non-polar cavities of FAPs, there has been an evolutionary shift to a higher prevalence of polar and positively charged residues among auxiliary CHILs and catalytic CHIs. Remarkably, Tyr^106^ was highly conserved across all type 1, 2, and 4 CHI-fold sequences. As expected, we observed high conservation of key residues associated with type 2 activity (i.e. Gly^95^, Lys^97^, Thr^190^, and Met^191^ based on the numbering of residues in MsCHI-II from *Medicago sativa*) across legume type 2 sequences. While legume type 1 sequences retained Met^191^, non-legumes exhibited an Ile at this position. Comparative CHIL sequence analysis revealed differences in the ligand-binding pockets between legumes and other angiosperms (eudicots and monocots). We noted that residues Tyr^48^, Leu^95^, and Ala^113^ in legumes were distinct from CHILs in other plant species, which were predominantly substituted with Thr/Asn, Val/Ile, and Ser/Ala, respectively.

**Fig. 2.**
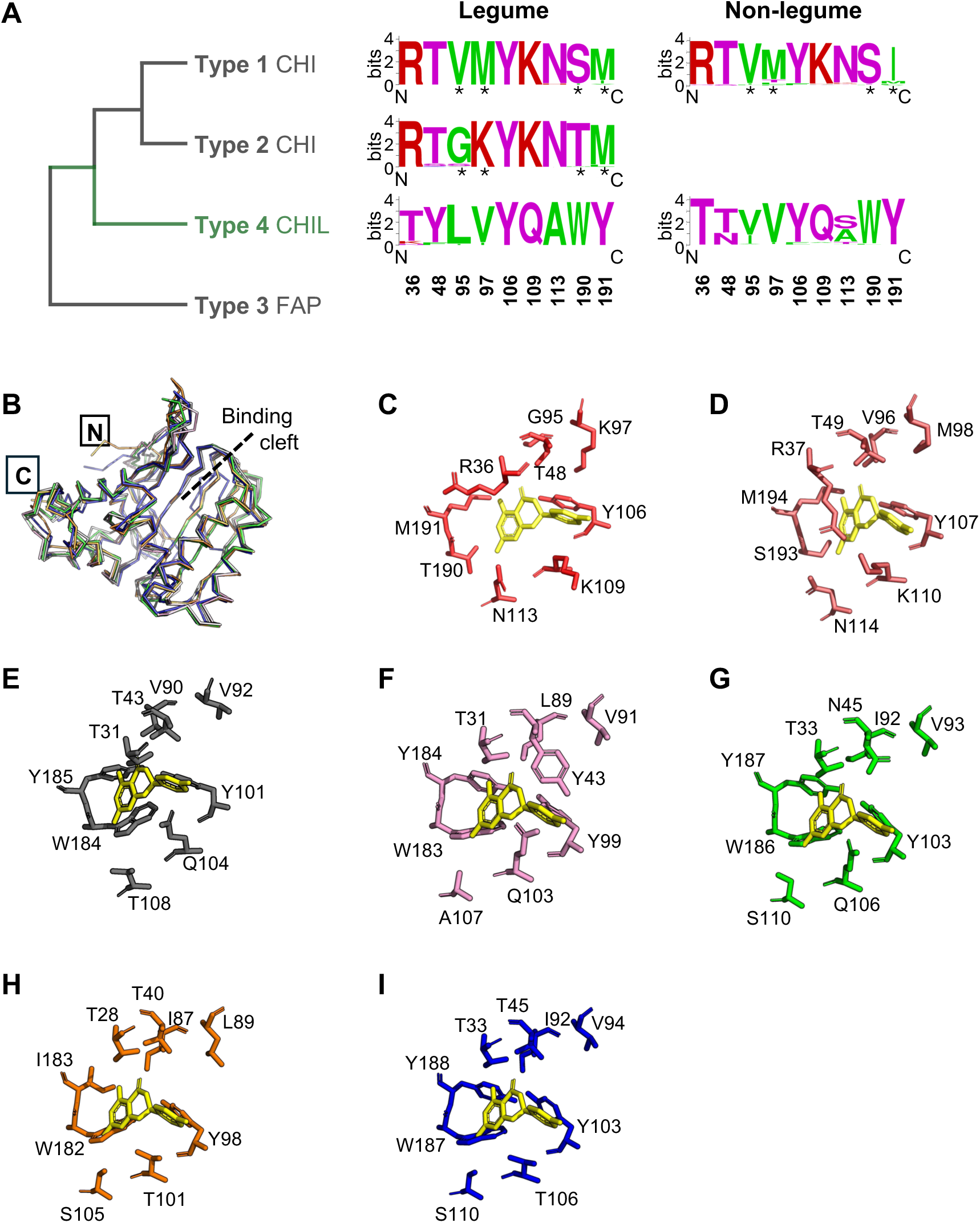
CHI and CHIL conserved binding cleft residues. **(A)** Sequence logos of extracted residues (36, 48, 95, 97, 106, 109, 113, 190, 191 based on MsCHI-II numbering) of type 1, 2, and 4 CHI-fold proteins in (non)-legumes. Residues are colour-coded: green, hydrophobic; purple, polar uncharged; red, positively charged. Asterisks indicate residues reported to be important for type 2 CHI behaviour. Basal bryophyte/lycophyte type 2 CHI sequences were not considered. **(B)** Structural alignment of the alpha carbon ribbon diagrams of GmCHIL (pink), OsCHIL (green), SmCHIL (orange), and MpCHIL (blue) models against AtCHIL resolved crystal structure (grey; PDB: 4DOK), **(C)** MsCHI-II (PBD: 1EYQ) structure and critical active site residues. **(D)** MtCHI-I (PDB: 6MS8) structure and critical active site residues. Residues of interest relative to bound NAR (based on MsCHI-II) for **(E)** AtCHIL, **(F)** GmCHIL, **(G)** OsCHIL, **(H)** SmCHIL, and **(I)** MpCHIL.

Following this survey, we selected a subset of four CHILs, sampling from across plant lineages. We chose CHIL sequence isoforms from *Glycine max* (GmCHIL; legume), *Oryza sativa* (OsCHIL; monocot), *Selaginella moellendorffii* (SmCHIL; lycophyte), and *Marchantia polymorpha* (MpCHIL; bryophyte). These four isoforms share similar primary structures, ranging from 205 to 229 amino acids in length, and share approximately 37-60% sequence identity (Supplementary Fig. 2). To facilitate further structural analysis, homology models of these four proteins were generated and aligned with the crystal structure of AtCHIL (PDB: 4DOK) (Fig. 2B). This structural comparison revealed a high degree of CHI-fold conservation, as indicated by the low root mean squared deviation (RMSD) values relative to AtCHIL: 0.745 (GmCHIL), 0.755 (OsCHIL), 0.943 (SmCHIL), and 1.057 Å (MpCHIL).

Interestingly, a comparison of CHIL binding clefts compared to the active sites of type 2 and type 1 CHIs, holo-MsCHI-II (PDB: 1EYQ; Fig. 2C) and holo-MtCHI-I (*M. truncatula*; PDB: 6MS8; Fig. 2D), respectively, uncovered key substitutions within the CHIL clefts that appear to restrict access significantly. A substrate NAR molecule was modelled in the respective binding cavities of each CHIL to investigate the interaction between these key residues and a prospective molecule (Fig. 2E-I). Compared to CHI, the binding clefts of CHILs exhibited several distinct structural differences. Notably, two bulky substitutions were identified: Thr/Ser^190^ in CHI was replaced by Trp in CHIL, while Met^191^ was replaced by Ile in SmCHIL or by Tyr in the remaining CHILs of interest. These substitutions may introduce steric hindrance into the cleft, altering interactions with and acting as significant obstructions to NAR accommodation. Additionally, the catalytic Lys^109^ in CHIs is substituted for Gln in the land plant isoforms studied here, which may further restrict ligand access to the binding site. In the GmCHIL cavity, the steric hindrance is expected to be further increased by Tyr^48^, which may particularly constrain the binding pockets of legume CHILs.

### CHIL orthologs differentially impede the lactone shunt

Purified recombinant GmCHIL, OsCHIL, SmCHIL, and MpCHIL were assayed in combination with GmCHS8 to assess their capacity for repressing the formation of CTAL and improving flux towards NC and NAR (Fig. 3A). In the assay, *p*-coumaroyl-CoA and malonyl-CoA were incubated with CHS alone or coupled to CHIL isoforms (4:1 molar excess of CHIL:CHS). Two peaks (*m*/*z* [271]^-^) were identified as NAR and CTAL (Fig. 3G). CHS alone resulted in 60.6% CTAL (percentage of total product), while all CHILs significantly reduced the accumulation of this by-product. GmCHIL provided the strongest suppression of CTAL formation to approximately 10% (Fig. 3D). The chalcone intermediate (NC) appears to have spontaneously isomerized to the flavanone product (NAR) during the incubation period. As indicated in previous characterizations of CHS, we did not perceive BNY in any of our reaction assays, likely due to its instability.

**Fig. 3.**
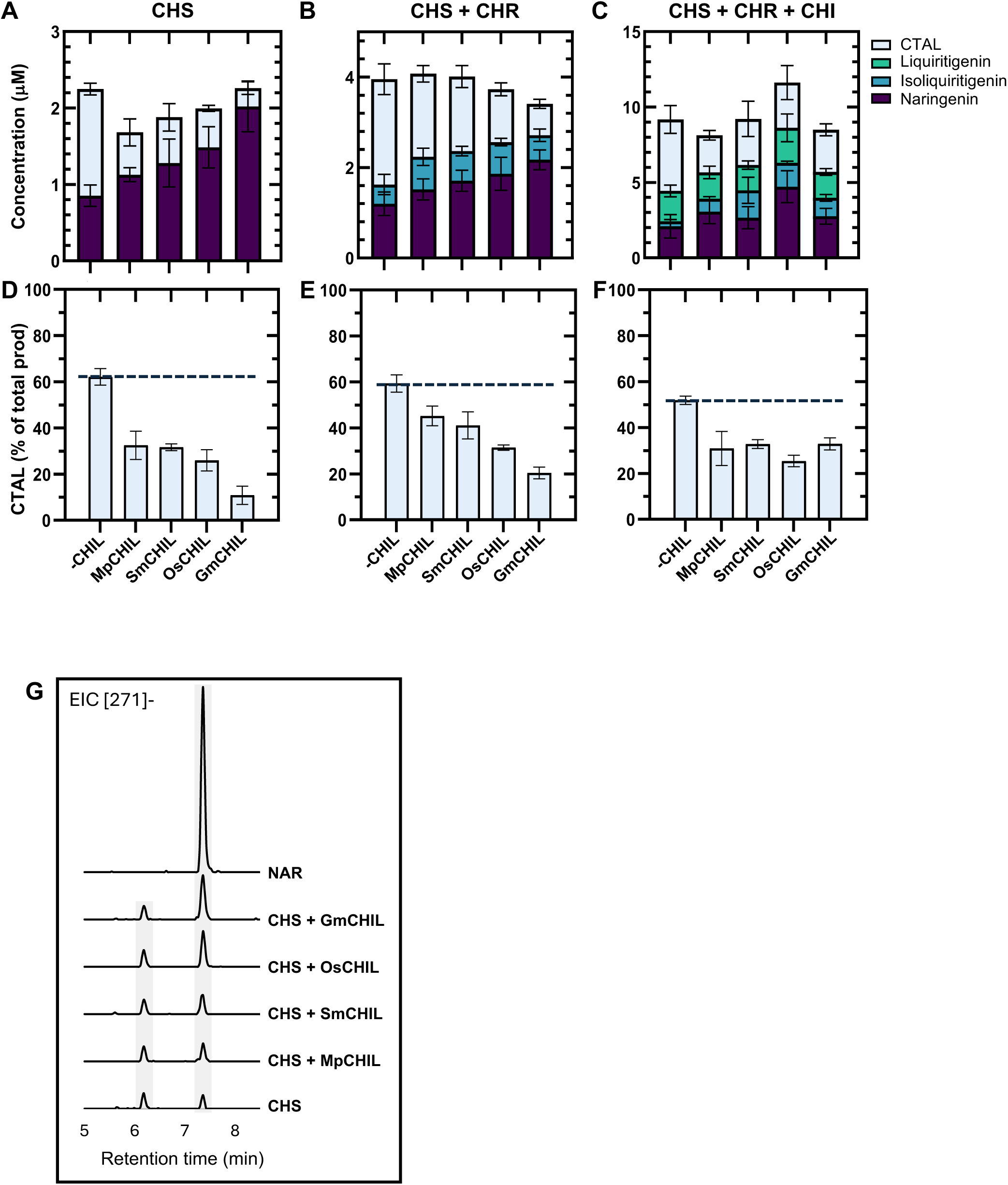
CHILs improve target compound yield by suppressing by-product formation *in vitro*. (A-C) Quantification of intermediates, products, and by-products from coupled-enzyme assays using CHS, CHR, and CHI. CHS was pre-incubated with or without CHIL for 10 min before adding 100 μM *p*-coumaroyl-CoA and 200 μM malonyl-CoA. **(D-F)** CHIL suppression of CTAL formation compared to total products measured. **(G)** Representative LC-MS extracted ion chromatograms for *m/z* [271]^-^. The peak at retention time (RT) 7.3 min corresponds to that of the NAR authentic standard, while the peak at RT 6.1 min represents the by-product CTAL.

The utility of CHIL in rectifying CHS promiscuity has been previously established; however, its impact on the ISO (6′-deoxychalcone) branch has not been determined. To access this branch, we supplemented CHS with GmCHR14A and NADPH, with or without CHIL. CHS-CHR coupled reactions produced three distinct peaks for CTAL, NAR, and ISO. CHS and CHR produced 59.94% by-product (Fig. 3B, E), which is not significantly different from the promiscuous activity exhibited by CHS alone. The inclusion of CHILs significantly reduced by-product formation to 43-21%, with GmCHIL exerting the most pronounced impact.

While NC spontaneously rearranges to NAR, the isomerization of ISO is inherently described as being blocked. The latter contains a high energy intramolecular C=O···H−O bond, which precludes significant occurrence of autocyclization, requiring a catalyst in the form of type 2 CHIs. We supplemented GmCHI1B2 to efficiently convert ISO to (*2S*)-LIQ, in addition to catalyzing the stereoselective formation of (*2S*)-NAR. Consistent with the previous assays, reactions with CHS, CHR, and CHI led to a significant excess of CTAL at 51.88 % of the total product (Fig. 3C, F). All CHIL isoforms significantly reduced by-product synthesis, approximately halving carbon flux through the aberrant lactone shunt.

### Yeast biotransformation of *p*-coumaric acid to NAR

For our engineered *Saccharomyces cerevisiae* strains, we employed a plasmid-based, galactose-inducible approach utilizing the pESC vector series. The pathway was extended one step upstream by incorporating 4-coumarate-CoA ligase (4CL; EC 6.2.1.12). This approach enabled us to start with *p-*coumaric acid as the feeding substrate, which is a more stable and cost-effective alternative to *p*-coumaroyl-CoA and malonyl-CoA. Biotransformation assays were conducted by co-expressing 4CL and CHS and incubating the strains with the substrate for 48 h, incorporating additional plasmids as indicated.

In the initial round of strain development, we selected *Gm4CL2* cloned from soybean leaf cDNA to provide 4CL function. The strain harbouring *pESC-His-CHS-Gm4CL2* (H1) exhibited strong expression of CHS but weak expression of Gm4CL2 (Fig. 4A). Notably, the putative band for FLAG-tagged Gm4CL2 (approximately 100 kDa versus expected size 62 kDa) was confirmed to be an artefactual immunoreactive endogenous yeast protein, as it was also present in uninduced and non-transformed BY4741ΔW strains (Supplementary Fig. 3). The H1 strain produced trace amounts of the final product, NAR, and no intermediates except for the by-product CTAL. As observed in the *in vitro* assays, it is assumed that our method does not detect *p*-coumaroyl-CoA and that NC spontaneously isomerizes to NAR. These results suggest that the weak expression of Gm4CL2 limits substrate conversion at the first step.

**Fig. 4.**
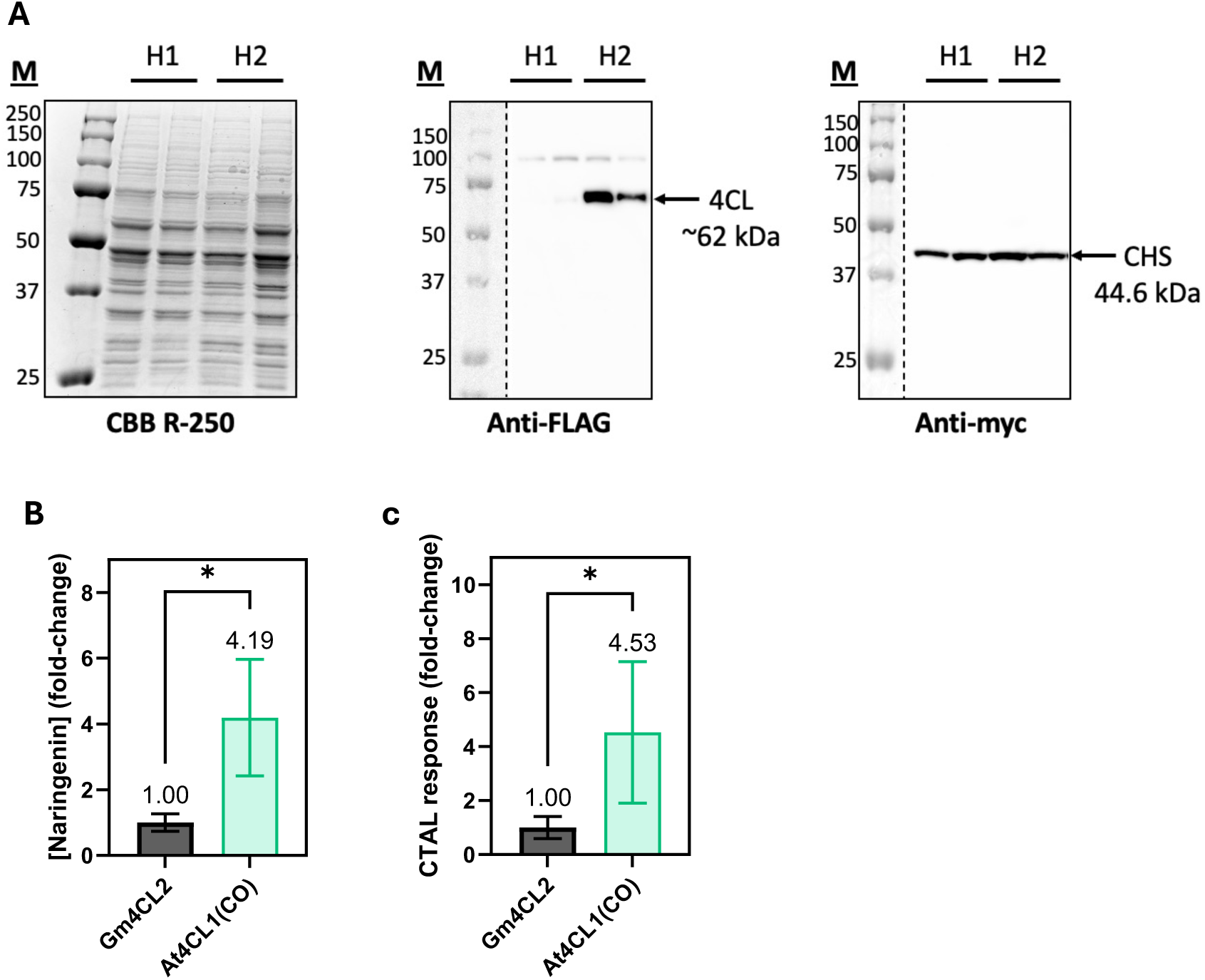
Increased 4CL expression leads to improved NAR titers. **(A)** Protein extracts of H1 and H2 yeast strains were subjected to SDS-PAGE and stained for total protein or immunoblotted against anti-FLAG or anti-myc. H1 strain harboured *pESC-Ura-CHS-Gm4CL2*, and H2 strain harboured *pESC-Ura-CHS-At4CL1(CO)*. Fold-change in **(B)** NAR titer and **(C)** CTAL response between H1 and H2 strains incubated with 250 μM *p*-coumaric acid for 48 h.

We replaced Gm4CL2 with a synthetic, codon-optimized At4CL1 to bypass this bottleneck. The new H2 strain co-expressing At4CL1 with CHS exhibited improved 4CL performance, considering both protein abundance and coupled activity; the latter was boosted approximately 4-fold (Fig. 4A, B). Lower rates of CTAL accumulation were observed in all yeast biotransformations compared to the *in vitro* assays, and were not quantifiable based on the NAR or NC calibration curves. However, CTAL levels did increase proportionally with NAR production in the H2 strain (Fig. 4B, C), indicating that the two 4CL orthologs do not differentially regulate CHS promiscuity.

### Synergistic effects of CHIL on isoflavonoid biosynthesis

H2 was further transformed with *pESC-Ura* encoding *CHI* alone (N1) or with one of *SmCHIL* (N2), *OsCHIL* (N3), or *GmCHIL* (N4). *MpCHIL* was not included in the remaining analyses as it demonstrated no expression in yeast as measured by immunoblots during preliminary trials. Furthermore, our *in vitro* analyses indicated minimal impact in promoting CHS fidelity compared to the remaining orthologs (Fig. 3).

The inclusion of CHILs increased NAR production by 25-60% (Fig. 5A). Immunoblotting crude soluble extracts of each strain after 48 h induction revealed strong expression of CHS and CHI and fair expression of 4CL, with no variation between strains (Supplementary Fig. 4B). However, expression levels of the CHILs were not uniform, with SmCHIL expressing the highest, followed by GmCHIL and OsCHIL. Therefore, the kinetic impacts of each CHIL on the pathway flux cannot be directly compared due to differences in stoichiometry. Despite the lower protein levels, strains containing OsCHIL (N3) and GmCHIL (N4) yielded the highest NAR titers (Fig. 5A). It is possible that increasing the stoichiometric ratio of OsCHIL and GmCHIL above other components could further enhance pathway flux. A minor peak (*m*/*z* [271]^-^) corresponding putatively to CTAL did not change significantly across the tested strains (Fig. 5D, G).

**Fig. 5.**
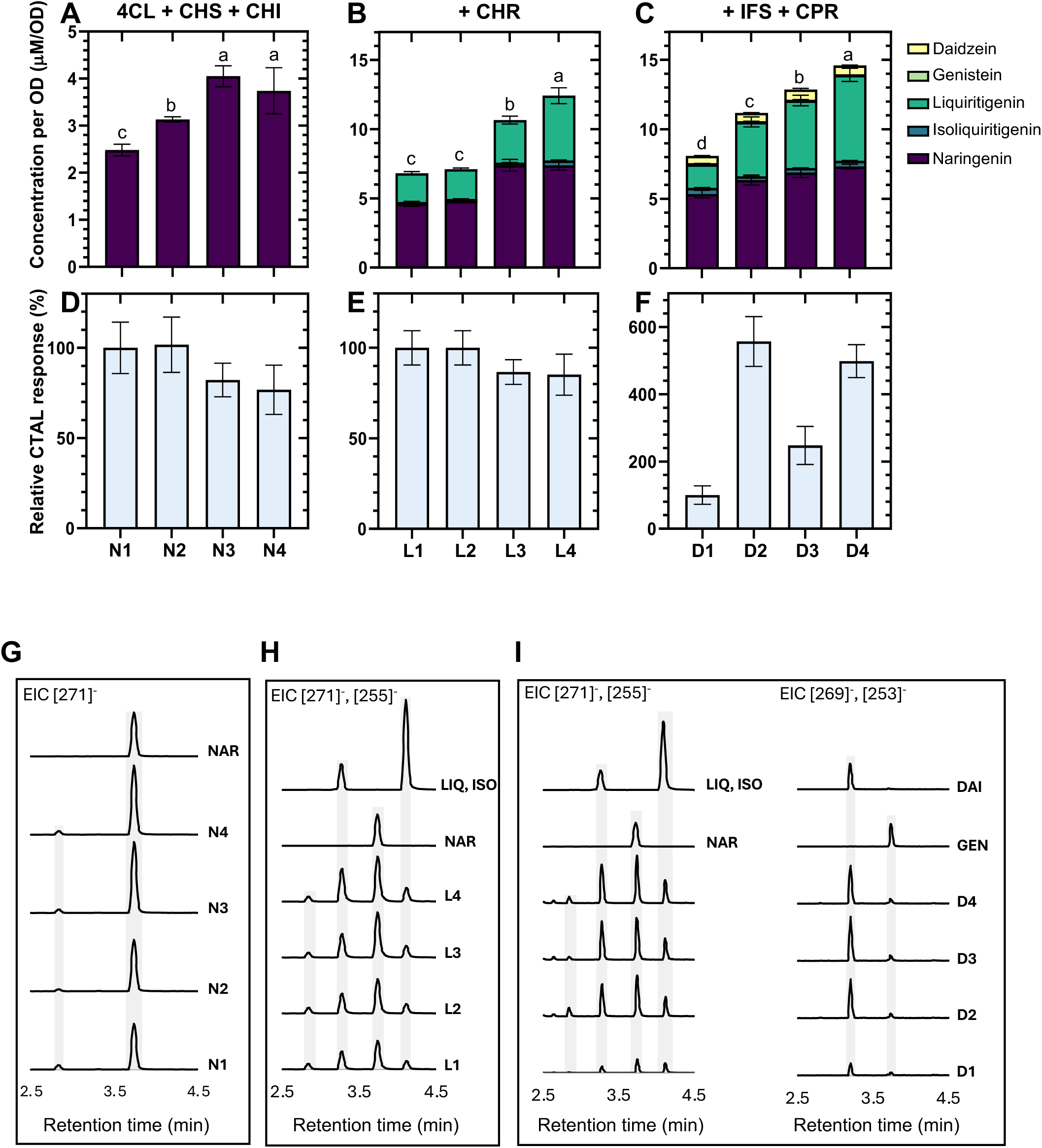
CHILs enhance pathway flux *in vivo*. **(A-C)** Quantified products and intermediates from yeast biotransformation assays. One-way ANOVA with Tukey’s HSD post-hoc test was performed on total products; letters indicate significant differences. **(D-F)** CTAL response rate was measured after 48 h biotransformation assays, wherein the response of strains lacking CHIL was set to 100% for each lineage (N, L, D). **(G-I)** LC/TQ-MS extracted ion chromatograms for *m*/*z* [271]^-^, [255]^-^, [269]^-^ and [255]^-^. Peaks were identified and quantified using retention times and multiple reaction monitoring (MRM) fragmentation patterns of authentic standards (NAR, LIQ, ISO, GEN and DZN). A minor peak with *m/z* [271]^-^ at RT 2.8 min was absent in the negative control reactions (without *-p*-coumaric acid substrate) and was identified as the by-product CTAL.

The H2 strain (At4CL; CHS) was also sequentially transformed with *pESC-Trp-CHR* to generate an L0 strain, followed by *pESC-Leu2d*-*IFS-CPR* to generate a D0 strain (Supplementary Fig. 4A). Both L0 and D0 strains were transformed with the aforementioned *pESC-Ura* constructs, yielding strains with CHI alone or CHI and one of the three CHILs.

The L-series strains (+CHR) produced a new peak corresponding to the 6ʹ-deoxychalcone, ISO, followed by the 5-deoxyflavanone, LIQ, with the incorporation of CHI (L1-4 strains). As expected, this type 2 CHI (GmCHI1B2) produced both flavanone outputs, NAR and LIQ (Fig. 5B, H). The partial diversion of chalcone intermediates into the deoxy-branch amounted to 20% of the total pathway flux in L0, with the addition of CHI (L1) or CHI with GmCHIL (L4) further enhancing this partitioning up to 30 and 40%, respectively (Figure 6A). Compared to the L1 control (no CHIL), NAR titers in both L3 and L4 improved by 60%, while LIQ titers in L3 and L4 increased by 50% and 125%, respectively (Fig. 5B and Supplementary Fig. 5A).

**Fig. 6.**
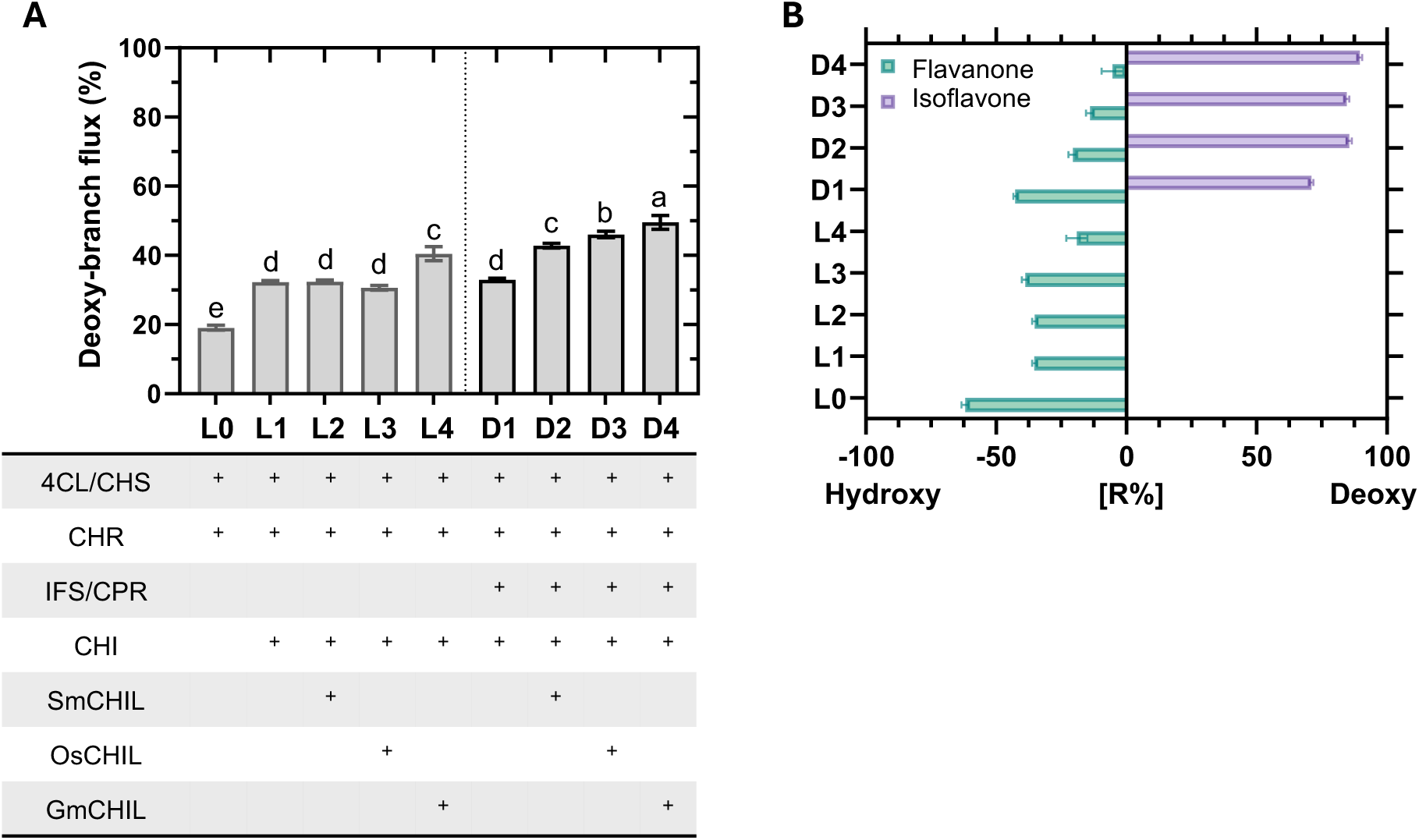
CHI-fold proteins mediate partitioning to the deoxy-branch. **(A)** The proportion of pathway flux into the deoxy-branch (CHR-catalyzed branch) for each strain. One-way ANOVA with Tukey’s HSD post-hoc test was performed; letters indicate significant differences. **(B)** CHR-branch dominance ([R%]) at both the flavanone and isoflavone steps for strains as indicated. [R%] was calculated as [(deoxy - hydroxy) ÷ (deoxy + hydroxy)] x 100.

Finally, the inclusion of GmIFS2 and a commonly employed plant P450 reductase partner, *Catharanthus roseus* CrCPR (D-series strains), gave rise to two new peaks corresponding to the isoflavones GEN and DZN (5-deoxyisoflavone) (Fig. 5I). IFS represents a rate-limiting step as all strains accumulated more flavanone intermediates compared to isoflavone products (Fig. 5C). The addition of SmCHIL (D2), OsCHIL (D3) and GmCHIL (D4) improved total intermediate/product accumulation by 38%, 58%, and 80%, respectively, compared to no CHIL (D1). This increase in total products is primarily attributed to increased NAR and LIQ titers rather than isoflavones. Nonetheless, all CHILs enhanced DZN production in these strains (D2-D4) by approximately 40-50%, whereas GEN production dropped by a similar percentage (Supplementary Fig. 5B).

While the deoxy-branch in D1 contributed to only ∼30% of the pathway flux, IFS displayed a 5-fold greater preference for the deoxy substrate (LIQ), yielding primarily DZN (Fig. 5C and Supplementary Fig. 5B). SmCHIL, OsCHIL, and GmCHIL had incrementally positive effects on flux shuttled through the deoxy-branch, which increased to 42%, 46%, and 50%, respectively (Fig. 6A). Further, CHR-branch dominance [R%] revealed CHIL-influenced diversion at the flavanone and isoflavone steps. The inclusion of CHIL brought the relative abundance of LIQ and NAR to a near-equilibrium, while the product selectivity for isoflavone production shifted closer to complete dominance of DZN production (Fig. 6B).

## DISCUSSION

In this study, we characterized the impact of auxiliary CHILs on the complete isoflavone biosynthetic pathway using an *in vitro* and *in vivo* approach. Beyond its known role in rectifying CHS promiscuity, we have uncovered an additional function of CHIL in regulating flux partitioning within the branched (iso)flavonoid pathway. By co-expressing CHIL with different combinations of the pathway components in yeast, we observed a remarkable synergistic effect of CHIL with IFS in driving metabolic flux through the deoxy-branch of the bifurcated pathway. These findings underscore the role of non-catalytic components in biosynthetic pathways and their potential in engineering metabolic content *ex planta*. It is mechanistically unclear how CHIL executes this function; however, previous research suggests protein-protein interactions with CHS and probable metabolon formation (21, 32).

CHILs are the only members of the CHI-fold that can rectify CHS promiscuity by inhibiting the lactone shunt. Here, as in previous research, we have explicitly shown that *bona fide* CHIs do not affect lactone (CTAL) accumulation (Fig. 3) (21). Furthermore, downstream enzymes (CHR and IFS) are also incapable of engendering CHS fidelity for chalcone production. The mechanism of CHIL function has been alluded to but has not been comprehensively described. Suggested possibilities include (i) an interaction of CHIL with CHS that would promote the canonical tetraketide formation and cyclization, (ii) the binding of intermediates after each round of acetylation, or (iii) the binding of the lactone by-products to inhibit their accumulation.

CHILs and CHIs appear to have evolved from a common ancestor associated with fatty acid biosynthesis, i.e., FAPs (16, 29). While the ligand binding site of FAPs encompasses a largely buried, non-polar cavity that sequesters the aliphatic chain(s) of fatty acids, the CHIs and CHILs possess more sterically restricted pockets with protruding polar residues (28). Although CHILs are non-catalytic and unable to rescue the *transparent testa* (*tt5*) *Atchi* mutant in Arabidopsis, a recent report suggests that catalytically impaired CHIs (type 1) could fulfill this role (21, 26, 27, 28, 33–34). Non-catalytic, T48A or Y106F mutant CHIs rescued the loss of flavonoid and anthocyanin content in *chi* loss-of-function mutants in *Zea mays* (maize) and *Solanum lycopersicum* (tomato). This unexpected finding suggests that the loss of critical active site residues might not completely abrogate chalcone-binding and that inactive CHIs potentially facilitate isomerization into the (*S*)-enantiomer for downstream enzymes, albeit at a much-reduced rate. Remarkably, Tyr^106^ is strongly conserved across all CHI-fold sequences, including non-catalytic CHILs, indicating significant selective pressure retained this residue for its structural or biochemical function (Fig. 2A). Moreover, structural comparisons reveal significant conservation of the CHI-fold between CHIs and CHILs, suggesting that CHIL may similarly bind chalcones or flavanones (28). However, the distinct architecture of the two suggests that CHIL could bind intermediates directly prior to chalcone formation (Fig. 2).

Molecular docking of NAR to the resolved *Vitis vinifera* (grapevine) VvCHIL structure suggests that bulky Gln^109^ and Trp^190^ residues may obstruct flavanone entry into the binding cleft (35). We noted other bulky substitutions in CHILs compared to holo-MsCHI-II and holo-MtCHI-I, including M191Y/I and T48Y in the case of legumes, which could contribute further to sterically restrict ligand entry or binding (Fig. 2). Ancestral reconstruction of a non-catalytic ancestor protein (ancCC), preceding CHIs and CHILs, suggested Arg^36^, Thr^48^, and Tyr^106^ were conserved binding cleft residues (29). Arg^36^ is critical for substrate alignment and stabilization in catalytic CHIs and non-catalytic ancCC and was inherited from FAPs, where it coordinates fatty acid carboxyl groups (28, 36). However, most CHIL sequences across diverse phyla have replaced Arg^36^ with a more compact and non-charged but polar Thr residue. These changes are expected to impact ligand binding in CHILs, including ligand preference, orientation and stabilization, and affinity. Therefore, it remains to be examined if CHILs preferentially bind the intermediates of CHS catalysis, such as the polyketide prior to cyclization, or the lactone by-products over the chalcone product.

The possibility of CHIL functioning as a ligand-binding partner or built-in biosensor for chalcone-flavanone biosynthesis has been explored, with some interesting outcomes. Thermal shift assays indicated a thermostabilizing effect of chalcone binding on various CHIL orthologs (35, 37). Indeed, CHILs from *V. vinifera* and *Physcomitrella patens* had a stronger affinity for chalcones (*k*_D_ ∼25-60 µM) than their flavanone counterparts (35). However, the authors suggest that such concentrations might not be physiologically relevant, questioning this putative role for CHILs. As chalcones cyclize spontaneously or enzymatically into flavanones *in planta*, NAR content can be used as a proxy for the cellular pool of these intermediates, with one study recording 2 µg/g NAR (FW) in *Malus domestica* (apple) leaves (38). Indeed, neither NAR nor NC are commonly detected in samples, as they are high-demand intermediates for many overlapping pathways. The apparent *K_m_* for NC in type 1 *Panicum virgatum* PvCHI and *Sorghum bicolor* SbCHI is ∼16 µM, while the apparent *K_m_* for ISO is 10 and 30 µM in type 2 GmCHI and MsCHI-II, respectively (16, 39, 40). Furthermore, the likelihood of CHIL serving as a chalcone-binding protein is diminished when considering that (i) chalcones serve as substrates with greater affinity for enzymes, including CHI and CHR (41), and (ii) CHIL-chalcone complexing does not explain how this protein suppresses by-product formation.

In contrast, CHIL-CHS interactions are well-documented and are thought to induce positive conformational changes in CHS that would enhance NC biosynthesis (21). While some interactions have been observed from pairs derived from diverse species, these interactions are preferentially conspecific (i.e., CHS and CHIL from the same species). Furthermore, CHILs from other species did not rescue the loss of proanthocyanidins in the *Arabidopsis thaliana* mutant *Atchil*, suggesting species-specific ortholog differences in binding affinity to AtCHS and function (15, 21). In contrast, our coupled assays demonstrated GmCHS rectifying activity in CHILs from diverse phyla, with the strongest outcomes pairing with GmCHIL, followed closely by OsCHIL (Fig. 3 and Fig. 5). These preferences are likely governed by interacting residues that have co-evolved in the two proteins in a species-dependent manner, although the exact molecular mechanism remains enigmatic.

While CHIL enhances CHS fidelity, an expanded NC pool alone does not fully explain its role in increasing deoxy-branch flux. Although NC was recently identified as the direct substrate for CHR, determining its kinetic parameters is challenging due to rapid spontaneous substrate turnover (41). Nonetheless, it is well established that CHIs are significantly more catalytically efficient than CHS and, by extension, CHR (21, 33, 39, 42). Moreover, type 2 CHIs favour NC over ISO, with GmCHI1B2 exhibiting over 7-fold higher catalytic efficiency with NC compared to the 6′-deoxychalcone (33). Consequently, an increased NC pool would be expected to favour flux towards NAR via CHI more than, or at least equally to, ISO via CHR. However, our data reveal that the presence of CHIL in the probable IFS-induced metabolon promotes CHR activity rather than driving CHI-catalyzed NC isomerization (Fig. 5). These findings suggest that CHIL may influence substrate channelling or modulate CHS-CHR interactions to activate deoxy-branch flux.

Auxiliary proteins, such as CHIL, are theorized to conduct further roles in metabolon function and flux partitioning (43, 44). In the isoflavonoid pathway, IFS is believed to be a nucleating member; it is anchored to the ER membrane, where it assembles soluble enzymatic partners on the cytosolic side (32, 45). Binary comparisons of our yeast L and D-series strains (the latter includes IFS) suggest a synergistic relationship between CHIL and IFS in promoting deoxy-branch utilization (Fig. 6). Indeed, it appears that IFS alone cannot drive flux through the deoxy-branch by complex formation with CHS-CHR. A catalytically nullified IFS (without a compatible CPR) was co-expressed with CHS, CHR, and CHI in yeast, but its presence unexpectedly reduced the deoxy-branch output moderately (46). The authors speculate that IFS is forming a complex; however, the lack of a compatible CPR in that system makes this assumption uncertain, as the complexing of P450-CPR has been shown to be necessary for metabolon formation (47, 48). Consequently, it is probable that while IFS is necessary for the aggregation of isoflavonoid enzymes to the ER, auxiliary proteins, such as CHIL, are responsible for regulating flux partitioning.

Enzyme promiscuity is a hallmark of many plant biosynthetic enzymes, driving the evolution of massive enzyme families with diverging substrate preferences and activities. This promiscuity has led to the expansion of plant specialized metabolism. However, it can also lead to loss of carbon flux into undesirable by-products, such as CTAL from CHS. While efforts to rationally engineer CHS have increased enzyme catalytic activity or broadened its polyketide product range, they have not succeeded in suppressing CTAL activity (49–51). Moreover, such strategies can have collateral and unexpected impacts on the coupled activity of enzymes in supramolecular complexes. In contrast, as an auxiliary protein, CHIL provides us with a straightforward solution to enhance CHS function in the bio-based production of valuable isoflavonoids. For instance, including CHIL could further enhance current yeast strains capable of *de novo* biosynthesis of DZN beyond the low mg/L range (52). Indeed, our results imply that CHILs play a more significant role in regulating (iso)flavonoid metabolism than previously understood, particularly in flux partitioning towards the deoxy-branch. Notably, DZN (5-deoxyisoflavone) is the precursor to many antimicrobials and pharmaceutically potent downstream compounds, including the pterocarpans. Therefore, including CHIL in bioengineering efforts will be crucial for accessing these valuable chemistries.

## MATERIALS AND METHODS

### Plant material

Soybean (*Glycine max* L. Merr. var DH410) was grown for six weeks with controlled conditions in growth chambers (16 hrs light, 8 hrs dark; 22 °C day, 20 °C night; 50-40 % humidity during day-night). They were fertilized once every 2 weeks (4 g/L Green Flag® all-purpose water-soluble fertilizer). Leaf tissues were harvested and flash-frozen in liquid nitrogen and stored at -80 °C.

### Chemicals

Analytical-grade standards were purchased for *in vitro* assays, including *p*-coumaric acid (Cayman Chemical, MI, USA), *p*-coumaroyl-CoA (MicroCombiChem, Wiesbaden, Germany), malonyl-CoA (Sigma Aldrich, MO, USA), naringenin chalcone (Phytolab, Lenexa, KS, USA), naringenin (Sigma Aldrich), isoliquiritigenin (Sigma Aldrich), liquiritigenin (Sigma Aldrich), genistein and daidzein (Cayman Chemical), NADPH (BioShop; Burlington, ON, CA), and fluorescein (Sigma Aldrich). Metabolites were stored at -20 °C in powder or lyophilized form.

### Multiple sequence alignment and homology modeling

The *Glycine max* genome was queried for representative type 1 (GmCHI2; Glyma.20G241700), type 2 (GmCHI1A; Glyma.20G241500), and type 4 (GmCHI4A; Glyma.06G143000) CHI-fold amino acid sequences from Phytozome v.13.0 by both keyword and sequence similarity searches. BLASTP was performed against all available genomes in Phytozome using these soybean sequences. The top 500 hits for each query were further processed to remove duplicate identifiers, ensuring a non-redundant dataset. The filtered sequences were aligned using Kalign and viewed in Jalview version 2.11.4.1 (53). Sequential logos were generated using WebLogo further to visualize trends in residues of interest (54).

AlphaFold Protein Structure Database (DeepMind) models were retrieved for GmCHIL (Q53B72), OsCHIL (Q2RBC7), SmCHIL (D8QX37), and MpCHIL (A0A2R6W2F7). The models were aligned to the reference structures for AtCHIL (PDB: 4DOK) and holo-MsCHI-II bound to naringenin (PDB: 1EYQ) in PyMol 3.1.1. The residues of interest identified in the multiple sequence alignments were visualized in relation to the naringenin molecule of the crystal MsCHI-II structure.

### RNA extraction and cDNA synthesis

Total RNA was isolated from approximately 70-100 mg of soybean leaf tissue and was homogenized using a Qiagen Tissue Lyser II® (Qiagen, Germany) with five 2.8 mm ceramic beads (OMNI International, GA, USA) per tube. Tissue homogenization was conducted in RLC buffer, followed by subsequent steps outlined in the Rneasy® Plant Mini Kit protocol (Qiagen), yielding 30 μl of the RNA sample. Total RNA was quantified using a NanoPhotometer® NP80 (Implen, CA, USA). Concentration and integrity were confirmed through gel electrophoresis of 0.2 μg of RNA on a 1% (w/v) agarose gel with 1% (v/v) bleach in 1x TAE buffer and stained with GelRed® (Sigma Aldrich) nucleic acid staining solution and Trackit™ CyanOrange loading buffer (Invitrogen). cDNA was synthesized from 1 μg of RNA using the All-In-One 5X RT MasterMix (ABM, BC, Canada), which includes genomic DNA removal, to a final concentration of 50 ng/μl. The cDNA thus prepared was the source for *Gm4CL2* and *GmIFS2* cloning.

### Cloning and microbial strains

The plasmids used in this study were obtained from the following sources. Plasmids used for recombinant protein purification in *E. coli*: *pET28α(+)-GmCHIL* (Azenta Life Sciences, MA, USA), *pET28α(+)-MpCHIL*, *pET28α(+)-SmCHIL*, and *pET28α(+)-OsCHIL* were produced synthetically and codon-optimized for *E. coli* expression (Twist Bioscience, CA, USA). The vectors containing soybean genes encoding biosynthetic enzymes of the pathway were acquired from Prof. Sangeeta Dhaubhadel, including *pET160-DEST-GmCHR14A*, *pET32α(+)-GmCHS8*, and *pDONR™/221-GmCHI1B2* (55, 56). Additionally, the following empty vectors were employed: *pET28α(+)* (Novagen, Germany), *pET160-DEST* (Invitrogen, MA, USA), and *pDONR™/Zeo* (Invitrogen). The *E. coli* strain NEB10*β* was used for plasmid DNA production (New England Biolabs (NEB); MA, USA), and BL21 (DE3) was used for recombinant protein expression.

To generate yeast strains, pESC yeast expression vectors (Agilent Technologies, CA, USA) were used to insert up to two genes of interest into the two multiple cloning sites (MCS) controlled by the reversely oriented promoters, *pGAL1* and *pGAL10*, using restriction digest cloning. Genes of interest included the same targets used for *E. coli* heterologous expression, with the addition of *Gm4CL2* and *GmIFS2,* which were cloned from soybean leaf cDNA, as well as synthetically generated *At4CL1*, codon-optimized for yeast expression (Twist Bioscience). All targets were C-terminally tagged, as indicated, and assembled into the following constructs: pESC-*His*-*GAL1:CHS-myc-GAL10:4CL-FLAG*, pESC-*Ura*-*GAL1:CHI-myc*, pESC-*Ura*-*GAL1:CHI-myc-GAL10:GmCHIL-FLAG*, pESC-*Ura*-*GAL1:CHI-myc-GAL10:OsCHIL-FLAG*, pESC-*Ura*-*GAL1:CHI-myc-GAL10:SmCHIL-FLAG*, pESC-*Ura*-*GAL1:CHI-myc-GAL10:MpCHIL-FLAG*, pESC-*Trp*-*GAL1:CHR-myc,* pESC-*Leu2d*-*GAL1:IFS-myc-GAL10:CrCPR-FLAG*. The latter construct was kindly provided by Prof. Yang Qu (University of New Brunswick) and features a cytochrome P450 reductase from *Catharanthus roseus*. Constructs were sequentially transformed into the *S. cerevisiae* BY4741ΔW strain, kindly provided by Prof. Ignea Codruta (McGill University) (Supplementary Fig. 4A and Supplementary Table 1).

### Recombinant protein production

*E. coli* BL21 (DE3) strains were cultured in LB at 37 °C and 225 rpm until reaching OD_600_ 0.5. Protein expression was induced by the addition of 1 mM isopropyl 1-thio-β-galactopyranoside (IPTG) (VWR International, Radnor, PA, USA) and further incubated at 25 °C for 4 h. Induced cells were harvested at 1,000 *x g* prior to resuspending (1:10 (w/v)) in buffer A (20 mM Tris-HCl pH 8.0, 300 mM NaCl, 10% glycerol (v/v)). Cells were sonicated on ice using a microprobe ultrasonic liquid processor (Misonix, NY, USA) at 60 Hz for 5 s on, 5 s off, for 5 min.

The crude soluble fraction was separated from cell debris by centrifuging at 14,000 *x g* for 30 min. His-tagged proteins were purified via batch/gravity-flow purification by incubating the clarified protein extracts with TALON® Superflow™ resin (Cytiva, Sweden) for 1 h at 4 °C with gentle agitation. The resin/sample mix was loaded onto a disposable polypropylene column (Thermo Fisher Scientific, MA, USA) and washed with 5 bed volumes of buffer A containing 5 mM imidazole. Target proteins were eluted using 10 bed volumes of buffer B (20 mM Tris-HCl pH 8.0, 300 mM NaCl, 150 mM imidazole, 10% glycerol (v/v)) and the final eluent was concentrated using Vivaspin® centrifugal filter units (10 kDa MWCO) (Sartorius, Germany). Protein concentrations were determined using the colorimetric Bradford protein assay dye (Bio-Rad, CA, USA) with bovine serum albumin standard (Bio-Rad, CA, USA). All centrifugation steps were conducted at 4 °C, and the final purified protein was flash-frozen and stored at -80 °C until usage.

### In vitro assays

*In vitro* assays were conducted in 50 μl reactions (50 mM HEPES-NaOH buffer pH 7.5, 100 μM *p-*coumaroyl-CoA, and 200 μM malonyl-CoA). Protein components included CHS (3 μg) alone or with the addition of a CHIL ortholog (4 μg). Denatured (boiled) CHIL proteins were used as negative controls. To assess the impact on the 6′-deoxychalcone branch or flavanone production, CHR (4 μg and 10 mM NADPH) and/or CHI (1 μg) were added, respectively. All reactions were pre-incubated without CHS at 30 °C for 10 min before the reaction was initiated with the addition of CHS and incubated for a further 40 minutes. Reactions were quenched with 20 μl of 1 N hydrochloric acid (HCl) and 180 μl of ethyl acetate and centrifuged at 16,000 x *g* for 30 min. The upper phase (100 μl) was dried down using the Savant SpeedVac™ DNA130 vacuum concentrator system (Thermo Fisher Scientific) and stored at -20 °C. Samples were resuspended in 80 μl methanol containing 500 μM fluorescein as an internal standard before running on the single-quadrupole LC-MS.

### Yeast protein induction, extraction, and microsomal preparation

Yeast strains were grown in 2 mL cultures overnight at 30 °C, 225 rpm, in SC media (2% glucose and appropriate dropout supplements) and were then subcultured into 500 µL of SC media containing 2% galactose as the sole carbon source (appropriate dropout supplements) in 96-deepwell microtitre plates to a final OD_600_ of 0.3. Following incubation for 48 h at 30 °C, 900 rpm, cells were harvested at 2,500 *x g* and washed with LTE buffer (20 mM Tris-HCl (pH 7.5), 200 mM LiOAc, and 2 mM EDTA) for 3 min, and 0.4 M NaOH for 5 min. Finally, cells were boiled in sodium dodecyl sulphate (SDS) sample buffer for 10 min prior to gel electrophoresis (see below).

To extract proteins associated with the microsomal fraction, 50 mL SC media (2% glucose and appropriate dropout supplements) cultures were grown for 24 h at 30 °C, 225 rpm. Cells were harvested and resuspended in fresh SC media containing 2% galactose as the sole carbon source. After 24 h incubation, cells were harvested and resuspended (1:3; w/v) in microsomal extraction buffer (25 mM HEPES (pH 6.8) containing 150 mM LiOAc, 250 mM sucrose, 1 mM phenylmethylsulfonyl fluoride (PMSF), and 5 mM *β*-mercaptoethanol) and glass beads added up to the meniscus. Cells were mechanically lysed by vortexing for 12 x 1 min intervals at 4 °C. Lysates were centrifuged at 10,000 *x g* for 2 min at 4 °C. The supernatant was further centrifuged at 21,000 *x g* for 1 h to pellet the crude microsome fraction. The resulting pellets were resuspended in microsomal extraction buffer before boiling them in SDS sample buffer to be immunoblotted.

### Gel electrophoresis and immunoblotting

All sodium dodecyl sulphate–polyacrylamide gel electrophoresis (SDS-PAGE) gels were composed of 4% (w/v) acrylamide stacking and 12% resolving gels. Electrophoresis was run at 200 V for at least 45 min or until the dye front ran off the gel. Gels were either stained for total protein using Coomassie Brilliant Blue R-250 (CBB R-250) or transferred onto poly(vinylidene) difluoride (PVDF) membrane for immunoblotting. Membranes were blocked in 5% (w/v) skim milk powder and incubated with mouse anti-FLAG or anti-myc (Sigma Aldrich) diluted 1:3,000 (v/v) in tris-buffered saline (pH 7.5) containing 0.1% tween (TBST), 1% (w/v) BSA, and 0.05% (w/v) NaN_3_. Immunoreactive bands probed with secondary goat anti-mouse antibody conjugated to horseradish peroxidase were treated with Clarity ECL substrates (Bio-Rad, CA, USA) and chemiluminescence emissions were detected by the ChemiDoc imager (Bio-Rad).

### Biotransformation assay and metabolite extraction

Yeast cultures were induced with 2% galactose at OD_600_ 0.3 in 96-deep-well plates and fed with 250 µM *p*-coumaric acid prior to incubation for 48 h at 30 °C, 900 rpm. Harvested cultures were treated with LTE buffer and then extracted with 75% ethanol. Following centrifugation at 21,000 *x g* for 30 min, the clarified supernatant was dried down and stored at -20 °C. To run extracts for mass spectrometry, samples were resuspended in 80% methanol containing 20 ng/mL fluorescein as an internal standard.

### LCMS

#### LC-Triple-Quadrupole/MS analysis; yeast biotransformation assays

Yeast biotransformation assay extracts were analyzed by the Agilent 6495 triple quadrupole LC/MS system (Agilent Technologies). An HPLC Poroshell 120 EC-C18 (3.0 x 50 mm, 2.7 μM) (Agilent Technologies) column was used for chromatographic separation at 30 °C with a guard column (Poroshell 120, UHPLC Guard.EC-C18, 3.0 mm). The binary solvent system was mobile phase A (H_2_O + 0.1% formic acid (FA)) and mobile phase B (acetonitrile (ACN) + 0.1% FA), using a flow rate of 0.4 ml/min. The column was equilibrated with 10% mobile phase B. Sample volume 1mL was injected, and the mobile phase B gradient was developed as follows: 10-30% B, 0-1 min; 10-50% 1-4 min; 50-90% 4-5 min; 90-100% 5-5.5 min. Both full scan (range of *m/z* 100-1000) and dynamic multiple reaction monitoring (dMRM) analyses were performed in negative mode using electrospray ionization. Quantification of intermediates was done in dMRM, filtering for the selected quantifier and qualifier ions of each compound at the respective retention times using authentic standards. CTAL response rate was identified based on the extracted ion chromatogram of [271]^-^ *m*/*z*.

#### LC-Single-Quadrupole/MS analysis; in vitro assays

*In vitro* assay products were measured using an Agilent 6120 LC-single-quadrupole/MS (Agilent Technologies). The following system was used to separate compounds: column (Poroshell 120 EC-C18 3.0 x 50 mm, 2.7 μM); guard (Poroshell 120, UHPLC Guard, EC-C18, 3.0 mm); flow rate of 0.5 ml/min; mobile phase A (H_2_O + 0.1% FA); mobile phase B, (ACN + 0.1% FA). The column was equilibrated with 95 % solvent A from 0-0.5 min. Subsequently, 5 μl of the samples were injected, and a gradient was developed by washing the column with 50 % solvent B (v/v) from 6- 17 min; 90 % (v/v) solvent B from 17-17.5 min, followed by 100 % solvent B from 17.5-19 min. The column was rewashed and equilibrated with 95 % solvent A. The full scan MS analysis was performed in negative ionization mode. The CTAL peak and corresponding retention time were confirmed by extracting the ion chromatogram for [271]^-^ *m*/*z*. Metabolites were detected (UV detector) using absorbance at 254 nm for fluorescein and LIQ, 280 nm for CTAL, NAR, and ISO, and 360 nm for NC. Calibration curves were generated for all compounds except CTAL, for which no authentic standard was available. Apparent CTAL concentrations were estimated using the NC calibration curve. Quantification was performed using the LC/MS OpenLab CDS software (Agilent Technologies).

## STATISTICS AND REPRODUCIBILITY

All experiments were conducted on a minimum of *n* = 3 biologically independent samples and analyzed by a two-tailed, unpaired Student’s *t*-test. To compare the significance of all groups within a set of assays, the data were analyzed using one-way analysis of variance (ANOVA) with subsequent pairwise comparisons conducted using Tukey’s HSD test (GraphPad Prism). Data is represented as sample mean ± standard deviation. Statistical significance was determined as *p<*0.05 and is denoted by letters. Letters denote statistically similar groups between and within treatments.

## DATA AVAILABILITY

The authors declare that the data supporting the findings of this study are available within the paper and its Supplementary Information files. Source data are provided within this paper.

## ACKNOWLEDGEMENTS

This work was funded by the Natural Science and Engineering Research Council of Canada (NSERC) Discovery Grant fund to MD (NSERC RGPIN-2021-02817) and the Centre de Recherche en Biologie Structurale (CRBS) studentship award to LMR. We thank colleagues for kindly providing constructs and strains. Dr. Sangeeta Dhaubhadel (Agriculture and Agri-Food Canada) gifted constructs encoding soybean enzymes (*pET160-DEST-GmCHR14A*, *pET32α(+)-GmCHS8*, and *pDONR™/221-GmCHI1B2*). Dr. Yang Qu (University of New Brunswick) gifted the pESC-*Leu2d*-*GAL10:CrCPR-FLAG* construct. Dr. Ignea Codruta (McGill University) gifted the *S. cerevisiae* strain BY4741ΔW. We thank Dr. Philippe Seguin for providing soybean seeds. Access to a single-quadrupole LC-MS was provided by Bastien Castagner (McGill University), and access to a triple-quadrupole was provided by the McGill Agilent Partnership Lab (MAPL).

## AUTHOR CONTRIBUTIONS

LMR and MD wrote the article. MS conducted phylogenetic analyses and generated the alignments with LMR, and LMR conducted subsequent protein homology modelling. LMR designed, conducted, and analyzed all yeast experiments. BS designed, conducted, and analyzed all *in vitro* enzyme assays. LMR conducted a final analysis of all data and generated the figures. SGL reviewed the manuscript. MD conceived the study, supervised the research, revised the manuscript, and approved it for submission.

