## Supplementary Figures for "Chalcone isomerase-like impedes the lactone shunt and enhances flux partitioning in a bifurcated pathway towards isoflavonoid biosynthesis"

### Supplementary Figure 1

#### Type 1

| Sequence | 36 | 48 | 95 | 97 | 106 | 109 | 113 | 190 | 191 |
|---|---|---|---|---|---|---|---|---|---|
| *MtCHI-I* | R | T | V | M | Y | K | N | S | M |
| *MsCHI-II* | R | T | G | K | Y | K | N | T | M |
| *Eucgr.J01153* | R | T | V | F | Y | K | N | - | - |
| *Eucgr.J01152* | R | T | V | F | Y | K | N | T | I |
| *Cocit.J0013* | R | T | V | F | Y | K | H | P | V |
| *AUR62027261* | - | - | - | - | Y | K | N | - | - |
| *AUR62015656* | R | T | - | - | Y | K | N | - | - |
| *Sapof.03G107200* | R | T | V | M | Y | N | M | S | I |
| *Bevul.2G108200* | R | T | I | M | Y | N | T | S | I |
| *EL10Ac2g03495* | R | T | I | M | Y | N | T | S | I |
| *Lus10030309.g* | - | - | - | M | Y | K | N | S | I |
| *Lus10003311.g* | R | T | V | M | Y | K | N | S | I |
| *Lus10030310.g* | R | T | V | M | Y | K | N | S | I |
| *evm_27.TU.AmTr_v1.0_scaffold00034.52* | R | T | V | M | F | K | K | S | I |
| *evm_27.TU.AmTr_v1.0_scaffold00034.53* | R | T | M | M | F | K | K | S | I |
| *Thupl.29381290s0001* | G | T | V | M | F | K | H | S | V |
| *Thupl.29347745s0001* | R | T | V | M | Y | K | N | S | I |
| *Ceric.33G035400* | R | T | V | F | Y | K | R | S | I |
| *151616* | R | T | V | M | F | K | N | T | M |
| *Dicom.08G043300* | R | T | V | L | Y | K | H | T | I |
| *Dicom.02G038100* | R | T | V | L | Y | K | H | T | I |
| *Zosma06g21110* | R | T | V | M | Y | K | N | S | I |
| *evm_27.TU.AmTr_v1.0_scaffold00034.116* | R | T | I | T | Y | K | N | S | I |
| *Nycol.F00121* | R | T | I | F | F | K | N | S | V |
| *Nycol.F00122* | R | T | I | L | Y | K | N | S | V |
| *Chala.04G052100* | R | T | V | L | Y | K | N | S | I |
| *Yucal.22G060800* | R | T | V | L | F | K | N | S | I |
| *AgateH2.22G057800* | R | T | V | L | F | K | N | S | I |
| *AgateH2.22G057900* | R | T | V | L | F | K | N | S | I |
| *evm.TU.AsparagusV1_07.317* | R | T | V | M | Y | K | N | S | I |
| *AgateH2.26G054500* | R | T | V | M | Y | K | N | S | I |
| *Yucal.26G052500* | R | T | V | M | Y | K | N | S | I |
| *YucfiAlt.26G058100* | R | T | V | M | Y | K | N | S | I |
| *AgateH2.02G167000* | R | T | V | M | Y | K | N | S | I |
| *YucfiAlt.02G266400* | R | T | V | M | Y | K | N | S | I |
| *Yucal.1Z193900* | R | T | V | M | Y | K | N | S | I |
| *Yucal.02G239700* | R | T | V | M | Y | K | N | S | I |
| *Dioal.09G071500* | R | T | V | M | Y | K | N | S | I |
| *EscalH2.2G227600* | R | T | I | M | Y | K | N | S | I |
| *Acora.07G100700* | R | T | V | M | Y | K | N | S | I |
| *Tylat.06G127700* | R | T | V | M | Y | K | N | S | M |
| *GSMUA_Achr4G16830_001* | R | T | V | L | Y | K | N | S | I |
| *Aco014232* | R | T | V | M | Y | K | N | S | I |
| *Joasc.10G103600* | R | T | V | M | Y | K | N | S | I |
| *Phala.01G033500* | R | T | I | M | Y | K | N | S | I |
| *OsKitaake03g392700* | R | T | V | M | Y | K | N | S | I |
| *HORVU5Hr1G112670* | R | T | V | M | Y | K | N | S | I |
| *Thint.S05G545700* | R | T | V | M | Y | K | N | S | I |
| *Traes_5BL_E86097AA2* | R | T | V | M | Y | K | N | S | I |
| *Thint.V05G479600* | R | T | V | M | Y | K | N | S | I |
| *Thint.J05G529700* | R | T | V | M | Y | K | N | S | I |
| *Brame.10PG131200* | R | T | V | M | Y | K | N | S | I |
| *Brame.02UG041000* | R | T | V | M | Y | K | N | S | I |
| *Barbu.2G364800* | R | T | V | M | Y | K | N | S | I |
| *Bradi1g03840* | R | T | V | M | Y | K | N | S | I |
| *Brahy.D01G0043800* | R | T | V | M | Y | K | N | S | I |
| *Brame.02PG173200* | R | T | V | M | Y | K | N | S | I |
| *Brahy.S02G0385600* | R | T | V | M | Y | K | N | S | I |
| *Brast02G359800* | R | T | V | M | Y | K | N | S | I |
| *Urofu.9G032800* | R | T | V | M | Y | K | N | S | I |
| *Pavir.9KG390224* | R | T | V | M | Y | K | N | S | I |
| *Pavir.5NG328900* | R | T | V | M | Y | K | N | S | I |
| *Pavir.9NG037000* | R | T | V | M | Y | K | N | S | I |
| *Sevir.9G033800* | R | T | V | M | Y | K | N | S | I |
| *Seita.9G034700* | R | T | V | M | Y | K | N | S | I |
| *Pahal.9G032500* | R | T | V | M | Y | K | N | S | I |
| *Chala.03G263300* | R | T | V | M | Y | K | N | S | I |
| *Chala.M006100* | R | T | V | M | Y | K | N | S | I |
| *Oropetium_20150105_23726* | R | T | V | M | Y | K | N | S | I |
| *ELECO.r07.3AG0210570* | R | T | V | T | Y | K | N | S | I |
| *ELECO.r07.3BG0255680* | R | T | V | M | Y | K | N | S | I |

| Sequence | 36 | 48 | 95 | 97 | 106 | 109 | 113 | 190 | 191 |
|---|---|---|---|---|---|---|---|---|---|
| *Pavag01G031900* | R | T | V | M | Y | K | N | S | I |
| *Pavag01G032000* | R | T | V | L | Y | K | N | S | I |
| *Zm00001d012972* | R | T | V | M | Y | K | N | S | I |
| *Zm00001d034635* | R | T | V | M | Y | K | N | S | I |
| *Misin02G012200* | R | T | V | M | Y | K | N | S | I |
| *Misin01G027800* | R | T | V | M | Y | K | N | S | I |
| *AndgeH2.01CG031200* | R | T | V | M | Y | K | N | S | I |
| *AndgeH2.01BG036800* | R | T | V | M | Y | K | N | S | I |
| *SoffiXsponR570.01Bg035600* | R | T | V | M | Y | K | N | S | I |
| *SoffiXsponR570.01Eg036900* | R | T | V | M | Y | K | N | S | I |
| *SoffiXsponR570.01Ag036300* | R | T | V | M | Y | K | N | S | I |
| *Sobic.001G035600* | R | T | V | M | Y | K | N | S | I |
| *SoffiXsponR570.09Bg149000* | R | T | V | M | Y | K | N | S | I |
| *SoffiXsponR570.09Cg159700* | R | T | V | M | Y | K | N | S | I |
| *SoffiXsponR570.09Dg118500* | R | T | V | M | Y | K | N | S | I |
| *SoffiXsponR570.9us90g075600* | R | T | V | M | Y | K | N | S | I |
| *SoffiXsponR570.09Ag159800* | R | T | V | M | Y | K | N | S | I |
| *SoffiXsponR570.5_9Ag404900* | R | T | V | M | Y | K | N | S | I |
| *SoffiXsponR570.04Cg255500* | R | T | V | M | Y | K | N | S | I |
| *SoffiXsponR570.01Dg006300* | R | T | V | M | Y | K | N | S | I |
| *Lalb_Chr06g0164121* | R | T | V | M | Y | K | N | S | M |
| *Medtr7g027135* | - | - | V | M | Y | K | N | I | I |
| *Sapur.010G169300* | R | T | V | M | Y | K | N | S | M |
| *Potri.010G213000* | R | T | V | M | Y | K | N | S | M |
| *Podel.10G218000* | R | T | V | M | Y | K | N | S | M |
| *VIT_213s0067g03820* | R | T | V | T | Y | K | N | S | I |
| *Ptrif.0004s0460* | R | T | V | M | Y | K | N | S | M |
| *Ciclev10032697m.g* | R | T | V | M | Y | K | N | S | M |
| *orange1.1g027531m.g* | R | T | V | M | Y | K | N | S | M |
| *Fxa7Cg101976* | R | T | V | M | Y | K | N | S | I |
| *FvH4_7g20870* | R | T | V | M | Y | K | N | S | I |
| *Fxa7Ag202088* | R | T | V | M | Y | K | N | S | I |
| *Fxa7Bg202027* | R | T | V | M | Y | K | N | S | I |
| *Fxa7Dg101819* | R | T | V | M | Y | K | N | S | I |
| *Prupe.2G225200* | R | T | V | T | Y | K | N | S | I |
| *MD07G1186300* | R | T | V | M | Y | K | N | S | I |
| *MD01G1118000* | R | T | V | T | Y | K | N | S | I |
| *MD01G1118100* | R | T | V | T | Y | K | N | S | I |
| *MD01G1117800* | R | T | V | T | Y | K | N | S | I |
| *CecanH2.2G032100* | R | T | V | M | Y | K | H | S | V |
| *CecanH2.1G004900* | R | T | V | M | Y | K | N | S | M |
| *CiPaw.01G280400* | R | T | V | M | Y | K | N | S | I |
| *Qurub.08G248300* | R | T | V | T | Y | K | N | S | I |
| *NdRogue1H2.08G206900* | R | T | V | T | Y | K | N | S | I |
| *Caden.06G217800* | R | T | V | T | Y | K | N | S | I |
| *BPChr06G09462* | R | T | V | M | Y | K | N | S | I |
| *Corav.Jeff.Hap2_g24547* | R | T | V | M | Y | K | N | S | I |
| *CamerWinkler.07G177300* | R | T | V | M | Y | K | N | S | I |
| *Kaladp0060s0328* | R | T | V | T | Y | K | N | S | I |
| *Kalax.1146s0001* | R | T | V | T | Y | K | N | S | I |
| *Kalax.0010s0201* | R | T | V | T | Y | K | N | S | I |
| *Aqcoe2G172600* | R | T | V | M | Y | K | N | S | I |
| *Sapof.03G107000* | R | T | V | M | Y | K | N | S | I |
| *FUN_004111* | R | T | V | M | Y | K | N | S | I |
| *AUR62015655* | R | T | V | M | Y | K | N | S | I |
| *AUR62020547* | R | T | V | M | Y | K | N | S | I |
| *Spov3_chr4.01127* | R | T | V | M | Y | K | N | S | I |
| *AH009939* | R | T | V | M | Y | K | L | - | - |
| *Bevul.2G107700* | R | T | V | M | Y | K | N | S | I |
| *EL10Ac2g03488* | R | T | V | M | Y | K | N | S | I |
| *LitulAlt.07G050600* | R | T | V | M | Y | K | N | S | I |
| *CKAN_00532200* | R | T | V | M | Y | K | N | S | M |
| *Lsat_1_v5_gn_9_66221* | R | T | V | M | Y | K | M | S | V |
| *HanXRQChr15g0496871* | R | T | V | M | Y | K | N | S | M |
| *HanXRQChr15g0496901* | R | S | V | L | F | K | N | S | Q |
| *HanXRQChr15g0496881* | R | T | V | M | F | K | N | S | M |
| *EhanaH2.03G119900* | R | T | V | M | Y | K | N | S | I |
| *Vadar_g17519* | R | T | V | M | Y | K | N | S | I |
| *Hyque.04G056200* | R | T | V | M | Y | K | N | S | M |
| *MyflaH2.07G045000* | R | T | V | M | Y | K | N | S | I |
| *Solyc05G000551* | R | T | V | M | Y | K | N | S | I |

**Supplementary Figure 1, contd.**

| Sequence | 36 | 48 | 95 | 97 | 106 | 109 | 113 | 190 | 191 |
|---|---|---|---|---|---|---|---|---|---|
| *Soltu.DM.05G001950* | R | T | V | M | Y | K | N | S | I |
| *Liphi.11G022900* | R | T | V | T | Y | K | N | S | I |
| *Migut.D00159* | R | T | V | T | Y | K | N | S | I |
| *HyleuH2.11G013600* | R | T | V | M | Y | K | N | S | V |
| *ChfasH2.4G105600* | R | T | V | M | Y | K | N | S | I |
| *ChfasH2.4G105500* | R | T | V | M | Y | K | N | S | I |
| *GlymaLee.20G202800* | R | T | V | M | Y | K | N | S | M |
| *GlysoPI483463.20G202800* | R | T | V | M | Y | K | N | S | M |
| *GlymaFiskIII.20G227000* | R | T | V | M | Y | K | N | S | M |
| *Glyma.20G241700* | R | T | V | M | Y | K | N | S | M |
| *Lj5g0014981* | R | T | V | M | Y | K | N | S | M |
| *Ca_18653* | R | T | V | M | Y | K | N | S | M |
| *Lalb_Chr02g0143921* | R | T | V | M | Y | K | N | S | M |
| *Medtr1g115890* | R | T | V | M | Y | K | N | S | M |
| *Medtr1g115870* | R | T | V | M | Y | K | N | S | M |
| *Tp57577_TGAC_v2_gene10384* | R | T | V | M | Y | K | N | S | M |
| *Vfaba.Hedin2.R1.Ung021880* | R | T | V | M | Y | K | N | S | M |
| *Ler.1DRT.1g084060* | R | T | V | M | Y | K | N | S | M |
| *Lcu.2RBY.1g077000* | R | T | V | M | Y | K | N | S | M |
| *arahy.Tifrunner.gnm1.ann1.VJQ7J1* | R | T | V | M | Y | K | N | S | M |
| *arahy.Tifrunner.gnm1.ann1.TJ3PHW* | R | T | V | M | Y | K | N | S | M |
| *Phcoc.07G008400* | R | T | V | M | Y | K | N | S | M |
| *Phacu.WLD.007G009000* | R | T | V | M | Y | K | N | S | M |
| *PvUI111.07G008500* | R | T | V | M | Y | K | N | S | M |
| *Phacu.WLD.002G307400* | R | T | V | M | Y | K | N | S | M |
| *Vigun07g288600* | R | T | V | M | Y | K | N | S | M |
| *Pl07G0000009400.v1* | R | T | V | M | Y | K | N | S | M |
| *Pl02G0000410700.v1* | R | T | V | M | Y | K | N | S | M |
| *PvUI111.02G271300* | R | T | V | M | Y | K | N | S | M |
| *Phcoc.02G272300* | R | T | V | M | Y | K | N | S | M |
| *MD07G1186400* | - | Y | V | F | Y | K | N | S | I |
| *Cucsa.182640* | R | T | V | L | Y | K | N | S | I |
| *DCAR_027694* | R | T | V | T | Y | K | N | S | I |
| *29740.t000017* | R | T | V | M | Y | K | N | - | - |
| *evm.TU.supercontig_1855.1* | R | T | V | M | Y | K | N | - | - |
| *Solyc05G000550* | R | S | V | L | F | K | N | S | M |
| *Soltu.DM.05G001960* | R | S | V | L | F | K | N | S | M |
| *Spipo29G0020600* | R | S | V | M | Y | K | N | T | I |
| *CAG027859* | R | T | V | M | Y | K | N | S | I |
| *evm.TU.Scaffold_2596.230* | R | T | V | M | Y | K | N | S | I |
| *CAG021505* | R | T | V | M | Y | K | N | S | I |
| *evm.TU.Scaffold_637.459* | R | T | V | M | Y | K | N | S | I |
| *Manes.07G107200* | R | T | V | T | Y | K | N | S | I |
| *Cocit.F1001* | R | T | V | M | Y | K | N | S | M |
| *Eucgr.F03816* | R | T | V | M | Y | K | N | S | M |
| *Thecc.10G061200* | - | - | L | K | Y | K | N | S | I |
| *Thecc.10G060600* | R | T | V | T | Y | K | N | S | I |
| *Gohir.D13G021000* | R | T | V | M | Y | K | N | S | I |
| *Gohir.A13G020400* | R | T | V | M | Y | K | N | S | I |
| *Anaoc.0016s1186* | R | T | V | M | Y | K | N | S | I |
| *Anaoc.0011s0700* | R | T | V | M | Y | K | N | S | I |
| *Clevi.0025s0581* | R | T | V | M | Y | K | N | S | I |
| *CsAcsn226.04G163400* | - | - | N | C | Y | K | N | S | I |
| *Ibeam.3236s0003* | R | T | V | M | Y | K | N | S | I |
| *Ibeam.0132s0044* | R | T | V | M | Y | K | N | S | I |
| *CsAcsn226.09G178700* | R | S | V | M | Y | R | N | S | F |
| *CsAcsn226.06G163900* | H | T | V | M | Y | R | N | L | F |
| *CsAcsn226.09G178300* | R | T | V | I | Y | R | N | L | F |
| *CsAcsn226.09G137600* | V | T | L | M | I | E | K | L | F |
| *CsAcsn226.09G178400* | Q | T | L | M | Y | G | K | S | F |
| *CsAcsn226.09G178600* | R | S | L | M | Y | G | K | S | F |
| *CsAcsn226.04G163300* | R | S | L | M | Y | G | K | - | - |
| *Bol018696* | R | T | V | M | Y | T | T | F | I |
| *Brara.K01350* | R | T | V | M | Y | I | T | F | I |
| *Sp5g07300* | R | T | V | M | Y | K | N | S | I |
| *Luann.0427s0040* | R | T | V | M | Y | K | N | S | I |
| *Thhalv10010658m.g* | R | T | V | M | Y | K | N | S | I |
| *Distr.0006s116400* | R | T | V | M | Y | K | N | S | I |
| *Eusyr.0007s0655* | R | T | V | M | Y | K | N | S | I |
| *Thlar.0021s0840* | R | T | V | M | Y | K | N | S | I |
| *Alyli.0198s0052* | R | T | V | M | Y | K | N | S | I |
| *Desop.0229s0191* | R | T | V | M | Y | K | N | S | I |
| *Alyli.0008s0233* | R | T | V | M | Y | K | N | S | I |
| *Crahi.0091s0056* | R | T | V | M | Y | K | N | S | I |
| *Isati.9424s0003* | R | T | V | M | Y | K | N | S | I |
| *Sialb.0006s0696* | R | T | V | M | Y | K | N | S | I |
| *Sialb.0201s0242* | R | T | V | M | Y | K | N | S | I |
| *Eruve.0527s0019* | R | T | V | M | Y | K | N | S | I |
| *Bol044343* | R | T | V | M | Y | K | N | S | I |
| *Camar.0201s0009* | R | T | V | M | Y | K | N | S | I |
| *Braju.09G374300* | R | T | V | M | Y | K | N | S | I |
| *Brara.I03775* | R | T | V | M | Y | K | N | S | I |
| *Brara.G01659* | R | T | V | M | Y | K | N | S | I |
| *Braju.07G160700* | R | T | V | M | Y | K | N | S | I |
| *Bol008652* | R | T | V | M | Y | K | N | S | I |
| *Braju.15G103000* | R | T | V | M | Y | K | N | S | I |
| *Sialb.0019s0144* | R | T | V | M | Y | K | N | S | I |
| *Sialb.0367s0078* | R | T | V | M | Y | K | N | S | I |
| *Sialb.1005s0004* | R | T | V | M | Y | K | N | S | I |
| *Crahi.0798s0009* | R | T | V | M | Y | K | N | S | I |
| *Myper.0005s0913* | R | T | V | M | Y | K | N | S | I |
| *Isati.3639s0006* | R | T | V | M | Y | K | N | S | I |
| *Isati.10103s0002* | R | T | V | M | Y | K | N | S | I |
| *Caamp.0095s0738* | R | T | V | M | Y | K | N | S | I |
| *Stapi.1952s0004* | - | - | V | M | Y | K | N | S | I |
| *Caamp.0078s0557* | R | T | V | M | Y | K | N | S | I |
| *Camar.2313s0007* | R | T | V | M | Y | K | N | S | I |
| *Eruve.0048s0014* | R | T | V | M | Y | K | N | S | I |
| *Braju.11G344400* | Q | T | V | M | Y | R | G | S | I |
| *Bol044344* | Q | T | V | M | Y | K | N | S | I |
| *Braju.09G374400* | Q | T | V | M | Y | K | N | S | I |
| *Brara.I03776* | Q | T | V | M | Y | K | N | - | - |
| *Mamar.0037s0511* | R | T | V | M | Y | K | N | S | I |
| *Bostr.0697s0129* | R | T | V | M | Y | K | N | S | I |
| *CsAcsn226.06G164000* | R | T | V | M | Y | K | N | S | I |
| *CsAcsn226.09G178500* | R | T | V | M | Y | K | N | S | I |
| *Carub.0005s2175* | R | T | V | M | Y | K | N | S | I |
| *Lesat.0112s0049* | R | T | V | M | Y | K | N | S | I |
| *AT3G55120* | R | T | V | M | Y | K | N | S | I |
| *AL5G36240* | R | T | V | M | Y | K | N | S | I |
| *Ah5G25440* | R | T | V | M | Y | K | N | S | I |
| *Lesat.0093s0008* | R | T | I | L | Y | K | N | S | I |
| *Roisl.0046s1508* | R | T | V | M | Y | K | N | S | I |
| *Roisl.0046s1507* | R | T | V | M | Y | K | N | S | I |
| *CsAcsn226.16G335100* | R | T | V | M | Y | K | N | S | I |
| *CsAcsn226.07G340300* | R | T | V | M | Y | K | N | S | I |
| *CsAcsn226.20G138200* | R | T | V | M | Y | K | N | S | I |
| *Alyli.0198s0051* | R | T | V | M | Y | K | N | S | I |
| *Desop.0229s0190* | R | T | V | M | Y | K | N | S | I |
| *Alyli.0008s0232* | R | T | V | M | Y | K | N | S | I |

### Supplementary Figure 1, contd.

#### Type 2

| | 36 | 48 | 95 | 97 | 106 | 109 | 113 | 190 | 191 |
|---|---|---|---|---|---|---|---|---|---|
| MsCHI-II | R | T | G | K | Y | K | N | T | M |
| CecanH2.2G031700 | R | T | G | K | Y | K | N | T | M |
| ChfasH2.4G105700 | R | T | G | K | Y | K | N | T | M |
| CecanH2.5G005000 | R | T | G | K | Y | K | N | T | M |
| CecanH2.5G005100 | R | T | G | K | Y | K | N | T | M |
| CecanH2.1G005000 | R | T | G | K | Y | K | N | T | M |
| arahy.Tifrunner.gnm1.ann1.2GDU51 | - | - | G | K | Y | K | N | T | M |
| Medtr1g115830 | R | E | G | K | Y | K | N | T | M |
| Phacu.WLD.003G248300 | - | - | G | K | Y | K | N | - | - |
| GlymaFiskIII.20G226800 | R | T | G | K | Y | K | N | T | M |
| GlymaLee.20G202600 | R | T | G | K | Y | K | N | T | M |
| GlysoPI483463.20G202600 | R | T | G | K | Y | K | N | T | M |
| Glyma.20G241500 | R | T | G | K | Y | K | N | T | M |
| Vigun07g288400 | R | T | G | K | Y | K | N | T | M |
| Pl07G0000009500.v1 | R | T | G | K | Y | K | N | T | M |
| Phacu.WLD.007G009100 | R | T | G | K | Y | K | N | T | M |
| Phcoc.07G008500 | R | T | G | K | Y | K | N | T | M |
| PvUI111.07G008600 | R | T | G | K | Y | K | N | T | M |
| Ca_18651 | R | T | G | K | Y | K | N | T | M |
| Tp57577_TGAC_v2_gene30361 | R | T | G | K | Y | K | N | T | M |
| Vfaba.Hedin2.R1.3g001120 | R | T | G | K | Y | K | N | T | M |
| Ler.1DRT.1g084020 | R | T | G | K | Y | K | N | T | M |
| Lcu.2RBY.1g076970 | R | T | G | K | Y | K | N | T | M |
| Medtr1g115820 | R | T | G | K | Y | K | N | T | M |
| arahy.Tifrunner.gnm1.ann1.425ZXH | R | T | G | K | Y | K | N | T | M |
| Lj5g0021768 | R | T | G | K | Y | K | N | T | M |
| Lj5g0007558 | R | T | G | K | Y | K | N | T | M |
| Lj5g0004759 | R | T | G | K | Y | K | N | T | M |
| Vigun07g288500 | R | T | G | K | Y | K | N | T | M |
| Phcoc.03G211700 | R | T | G | K | Y | K | N | T | M |
| PvUI111.03G215100 | R | T | G | K | Y | K | N | T | M |
| Phacu.WLD.003G248000 | R | T | G | K | Y | K | N | T | M |
| Pl03G0000319700.v1 | R | T | G | K | Y | K | N | T | M |
| Ler.1DRT.1g084040 | R | T | G | K | Y | K | N | T | M |
| Lcu.2RBY.1g076980 | R | T | G | K | Y | K | N | T | M |
| GlymaLee.20G202700 | R | T | G | K | Y | K | N | T | M |
| GlymaFiskIII.20G226900 | R | T | G | K | Y | K | N | T | M |
| GlysoPI483463.20G202700 | R | T | G | K | Y | K | N | T | M |
| Glyma.20G241600 | R | T | G | K | Y | K | N | T | M |
| GlysoPI483463.10G248800 | R | T | G | K | Y | K | N | T | M |
| Glyma.10G292200 | R | T | G | K | Y | K | N | T | M |
| GlymaFiskIII.10G277100 | R | T | G | K | Y | K | N | T | M |
| GlymaLee.10G251200 | R | T | G | K | Y | K | N | T | M |
| arahy.Tifrunner.gnm1.ann1.LGAM8W | R | T | G | K | Y | K | N | - | - |
| arahy.Tifrunner.gnm1.ann1.6JHV2K | R | T | G | K | Y | K | N | - | - |
| arahy.Tifrunner.gnm1.ann1.TY7N4H | R | S | G | K | Y | K | N | - | - |
| arahy.Tifrunner.gnm1.ann1.9C11CW | R | S | G | K | Y | K | N | T | L |
| arahy.Tifrunner.gnm1.ann1.9IBE58 | R | S | G | K | Y | K | N | T | L |
| arahy.Tifrunner.gnm1.ann1.Q5CKGN | R | S | G | K | Y | K | N | T | L |
| Vfaba.Hedin2.R1.3g001040 | R | T | S | K | Y | K | N | T | M |
| Ler.1DRT.1g084050 | R | T | S | K | Y | K | N | T | M |
| Lcu.2RBY.1g076990 | R | T | S | K | Y | K | N | T | M |
| Ca_18652 | R | T | G | K | Y | K | N | T | M |
| Medtr1g115850 | R | T | C | K | Y | K | N | T | M |
| Medtr1g115840 | R | T | S | K | Y | K | N | T | M |
| Tp57577_TGAC_v2_gene10424 | R | T | G | K | Y | K | N | T | M |

### Supplementary Figure 1, contd.

#### Type 4

| | 36 | 48 | 95 | 97 | 106 | 109 | 113 | 190 | 191 |
|---|---|---|---|---|---|---|---|---|---|
| *MtCHI-I* | R | T | V | M | Y | K | N | S | M |
| *MsCHI-II* | R | T | G | K | Y | K | N | T | M |
| *AtCHIL* | T | T | V | V | Y | Q | T | W | Y |
| *YucfiAlt.22G069400* | S | K | F | I | E | K | K | I | Y |
| *Mapoly0175s0004* | T | T | I | V | Y | T | S | W | Y |
| *227414* | T | T | I | L | Y | T | S | W | I |
| *Dicom.15G035000* | T | T | V | F | Y | L | S | W | I |
| *Sphfalx16G077000* | T | T | V | I | Y | S | S | V | Y |
| *Sphmag16G076700* | T | T | V | I | Y | S | S | V | Y |
| *CepurR40.1G175200* | T | T | I | I | Y | A | S | L | Y |
| *Pp6c4_13620* | T | T | I | I | Y | A | S | L | Y |
| *Pp3c4_25770* | T | T | I | I | Y | A | S | L | Y |
| *Pp6c26_2490* | T | T | I | I | Y | A | S | L | Y |
| *Pp3c26_4040* | T | T | I | I | Y | A | S | L | Y |
| *Ceric.09G016600* | Y | Y | I | I | F | P | T | W | L |
| *Thint.J05G181600* | T | N | V | V | Y | Q | S | W | Y |
| *Thint.S05G216600* | T | N | V | V | Y | Q | S | W | Y |
| *Traes_5BL_4F4E0B862* | T | N | V | V | Y | Q | S | W | Y |
| *Thint.V05G161400* | T | N | V | V | Y | Q | S | W | Y |
| *HORVU5Hr1G046480* | T | N | V | V | Y | Q | S | W | Y |
| *Brame.05UG401500* | T | N | V | V | Y | Q | S | W | Y |
| *Brame.05PG013700* | T | N | V | V | Y | Q | S | W | Y |
| *Brast05G291700* | T | N | V | V | Y | Q | S | W | Y |
| *Brahy.S05G0319100* | T | N | V | V | Y | Q | S | W | Y |
| *Barbu.5G540700* | T | N | V | V | Y | Q | S | W | Y |
| *Brasyl.5G511700* | T | N | V | V | Y | Q | S | W | Y |
| *Bradi4g44390* | T | N | V | V | Y | Q | S | W | Y |
| *Brahy.D04G0637400* | T | N | V | V | Y | Q | S | W | Y |
| *Phala.09G012100* | T | N | I | V | Y | Q | S | W | Y |
| *OsKitaake11g008100* | T | N | I | V | Y | Q | S | W | Y |
| *LOC_Os11g02440* | T | N | I | V | Y | Q | S | W | Y |
| *OsKitaake12g011800* | T | N | I | V | Y | Q | S | W | Y |
| *LOC_Os12g02370* | T | N | I | V | Y | Q | S | W | Y |
| *ELECO.r07.9AG0672530* | T | T | V | V | Y | Q | S | W | Y |
| *ELECO.r07.9BG0696100* | T | T | V | V | Y | Q | S | W | Y |
| *Oropetium_20150105_24750* | T | N | V | V | Y | Q | S | W | Y |
| *Oropetium_20150105_22136* | T | N | V | V | Y | Q | S | W | Y |
| *Chala.09G182500* | T | N | V | V | Y | Q | S | W | Y |
| *Chala.12G021600* | T | N | V | V | Y | Q | S | W | Y |
| *Pavir.4KG345500* | S | P | - | V | Y | Q | S | W | Y |
| *Pahal.8G008800* | T | N | V | V | Y | Q | S | W | Y |
| *Pahal.3G008000* | T | N | V | V | Y | Q | S | W | Y |
| *Pavir.8NG149800* | T | N | V | V | Y | Q | S | W | Y |
| *Pavir.3KG041700* | T | N | V | V | Y | Q | S | W | Y |
| *Pavir.8KG022400* | T | N | V | V | Y | Q | S | W | Y |
| *Sevir.7G312700* | T | N | V | V | Y | Q | S | W | Y |
| *Seita.7G301600* | T | N | V | V | Y | Q | S | W | Y |
| *Sevir.8G012200* | T | N | V | V | Y | Q | S | W | Y |
| *Seita.8G013300* | T | N | V | V | Y | Q | S | W | Y |
| *Urofu.3G033800* | T | N | V | V | Y | Q | S | W | Y |
| *Urofu.8G005700* | T | N | V | V | Y | Q | S | W | Y |
| *Pavag08G019100* | T | N | V | V | Y | Q | A | W | Y |
| *Zm00001d044683* | T | N | V | V | Y | Q | S | W | Y |
| *AndgeH2.08BG026900* | T | N | V | V | Y | Q | S | W | Y |
| *AndgeH2.08CG031600* | T | N | V | V | Y | Q | S | W | Y |
| *Sobic.008G030100* | T | N | V | V | Y | Q | S | W | Y |
| *AndgeH2.08AG030300* | T | N | V | V | Y | Q | S | W | Y |
| *Misin14G043700* | T | N | V | V | Y | Q | S | W | Y |
| *SoffiXsponR570.6_9Ag038300* | T | N | V | V | Y | Q | S | W | Y |
| *Misin15G023200* | T | N | V | V | Y | Q | S | W | Y |
| *Misin10G113900* | T | N | V | V | Y | Q | S | W | Y |
| *SoffiXsponR570.6us88g110900* | T | N | V | V | Y | Q | S | W | Y |
| *SoffiXsponR570.09Ag037000* | T | N | V | V | Y | Q | S | W | Y |
| *SoffiXsponR570.09Eg034500* | T | N | V | V | Y | Q | S | W | Y |
| *SoffiXsponR570.09Bg036400* | T | N | V | V | Y | Q | S | W | Y |
| *SoffiXsponR570.9os1g003500* | T | N | V | V | Y | Q | S | W | Y |
| *SoffiXsponR570.09Cg030400* | T | N | V | V | Y | Q | S | W | Y |
| *Thupl.29379046s0007* | T | T | V | I | Y | Q | A | W | Y |
| *evm.TU.AsparagusV1_02.366* | T | N | V | V | F | Q | A | W | Y |
| *YucfiAlt.18G068200* | T | N | V | V | F | Q | A | W | Y |

| | 36 | 48 | 95 | 97 | 106 | 109 | 113 | 190 | 191 |
|---|---|---|---|---|---|---|---|---|---|
| *AgateH2.18G066300* | T | N | V | V | F | Q | A | W | Y |
| *Yucal.1Z190600* | R | N | V | V | F | Q | A | W | Y |
| *Yucal.05G034400* | - | - | V | V | F | Q | A | W | Y |
| *evm.TU.AsparagusV1_07.1354* | T | N | F | V | - | - | - | - | - |
| *GSMUA_Achr11G23630_001* | T | N | I | V | Y | Q | A | W | Y |
| *Dioal.19G120300* | T | N | V | V | Y | Q | A | W | Y |
| *evm_27.TU.AmTr_v1.0_scaffold00077.54* | T | T | I | V | Y | Q | A | W | Y |
| *Tylat.06G115500* | T | N | I | V | Y | Q | S | W | Y |
| *Aco012547* | T | N | V | V | Y | Q | S | W | Y |
| *Tylat.12G057300* | T | N | I | V | Y | Q | S | W | Y |
| *Joasc.14G112300* | T | N | I | V | Y | Q | A | W | Y |
| *Joasc.14G112400* | T | N | I | V | Y | Q | A | W | Y |
| *LitulAlt.15G009400* | T | T | - | - | - | - | - | - | - |
| *EscalH2.1G455800* | T | T | V | V | Y | Q | W | W | Y |
| *Kaladp0074s0080* | T | T | V | V | Y | Q | A | W | Y |
| *Kalax.0086s0049* | T | T | V | V | Y | Q | A | W | Y |
| *Kalax.0565s0024* | T | T | V | V | Y | Q | A | W | Y |
| *Nycol.J01677* | T | N | I | V | Y | Q | A | W | Y |
| *Clevi.0011s1093* | T | T | V | V | Y | Q | S | W | Y |
| *Roisl.0042s0137* | T | T | V | V | Y | Q | A | W | Y |
| *Luann.0050s0072* | T | T | V | V | Y | Q | A | W | Y |
| *Crahi.0001s0207* | T | T | V | V | Y | Q | T | W | Y |
| *AT5G05270* | T | T | V | V | Y | Q | T | W | Y |
| *Ah6G05290* | T | T | V | V | Y | Q | S | W | Y |
| *AL6G15110* | T | T | V | V | Y | Q | T | W | Y |
| *Sp6g37560* | T | T | V | V | Y | Q | A | W | Y |
| *Stapi.8357s0001* | T | T | V | V | Y | Q | A | W | Y |
| *Caamp.0105s1602* | T | T | V | V | Y | Q | A | W | Y |
| *Camar.0041s0002* | T | T | V | V | Y | Q | A | W | Y |
| *Camar.8960s0001* | T | T | V | V | Y | Q | A | W | Y |
| *Bol044046* | T | T | V | V | Y | Q | S | W | Y |
| *Brara.J02674* | T | T | V | V | Y | Q | S | W | Y |
| *Ibeam.4335s0006* | T | T | V | V | Y | Q | S | W | Y |
| *Ibeam.11562s0001* | T | T | V | V | Y | Q | A | W | Y |
| *Thlar.0026s0247* | T | T | V | V | Y | Q | S | W | Y |
| *Lesat.0131s0130* | T | T | V | V | Y | Q | A | W | Y |
| *Lesat.0165s0477* | T | T | V | V | Y | Q | A | W | Y |
| *Eruve.0004s0108* | T | T | V | V | Y | Q | S | W | Y |
| *Sialb.0029s0048* | T | T | V | V | Y | Q | T | W | Y |
| *Braju.10G256100* | T | T | V | V | Y | Q | S | W | Y |
| *Sialb.0072s0169* | T | T | V | V | Y | Q | T | W | Y |
| *Sialb.0929s0036* | T | T | V | V | Y | Q | T | W | Y |
| *Myper.0004s0666* | T | T | V | V | Y | Q | A | W | Y |
| *Isati.1532s0002* | T | T | V | V | Y | Q | A | W | Y |
| *Isati.6513s0007* | T | T | V | V | Y | Q | A | W | Y |
| *Isati.0492s0025* | T | T | V | V | Y | Q | A | W | Y |
| *Isati.1396s0008* | T | T | V | V | Y | Q | A | W | Y |
| *Isati.0586s0031* | T | T | V | V | Y | Q | A | - | - |
| *Caamp.1038s0815* | T | T | V | V | Y | Q | A | W | Y |
| *Thhalv10014675m.g* | T | T | V | V | Y | Q | A | W | Y |
| *Braju.16G313600* | T | T | V | V | Y | Q | A | W | Y |
| *Bostr.13129s0438* | T | T | V | V | Y | Q | A | W | Y |
| *Carub.0006s0420* | T | T | V | V | Y | Q | A | W | Y |
| *Distr.0002s37500* | T | T | V | V | Y | Q | A | W | Y |
| *Eusyr.0039s0043* | T | T | V | V | Y | Q | A | W | Y |
| *CsAcsn226.20G040300* | T | T | V | V | Y | Q | S | W | Y |
| *CsAcsn226.13G041800* | T | T | V | V | Y | Q | T | W | Y |
| *Mamar.0004s0424* | T | T | V | V | Y | Q | T | W | Y |
| *CsAcsn226.08G286200* | T | T | V | V | Y | Q | T | W | Y |
| *Alyli.0192s0014* | T | T | V | V | Y | Q | T | W | Y |
| *Alyli.0180s0030* | T | T | V | V | Y | Q | T | W | Y |
| *Desop.0094s0101* | T | T | V | V | Y | Q | T | W | Y |
| *Lsat_1_v5_gn_6_40701* | T | T | I | V | Y | E | A | W | Y |
| *HanXRQChr08g0221541* | T | T | I | A | Y | Q | S | W | Y |
| *Lsat_1_v5_gn_8_11480* | T | T | I | V | Y | Q | A | W | Y |
| *Hyque.07G131600* | T | T | I | V | Y | Q | A | W | Y |
| *Hyque.04G090300* | T | T | I | V | Y | Q | A | W | Y |
| *Soltu.DM.05G022280* | T | T | V | V | Y | Q | A | W | Y |
| *Solyc05G002346* | T | T | V | V | Y | Q | A | W | Y |
| *EhanaH2.03G009800* | T | T | V | V | Y | Q | A | W | Y |

**Supplementary Figure 1, contd.**

| | 36 | 48 | 95 | 97 | 106 | 109 | 113 | 190 | 191 |
|---|---|---|---|---|---|---|---|---|---|
| *EhanaH2.02G133000* | T | T | V | V | Y | Q | A | W | Y |
| *Oeu044164.1* | T | T | V | V | Y | Q | A | W | Y |
| *Liphi.01G228500* | T | T | V | V | F | Q | A | W | Y |
| *Migut.F01245* | T | T | V | V | F | Q | A | W | Y |
| *evm.TU.Scaffold_637.822* | T | T | I | V | Y | Q | A | W | Y |
| *CAG021083* | T | T | I | V | Y | Q | A | W | Y |
| *CAG027433* | T | T | I | V | Y | Q | A | W | Y |
| *evm.TU.Scaffold_2596.579* | T | T | I | V | Y | Q | A | W | Y |
| *Acora.10G016000* | T | T | I | Q | Y | Q | A | W | Y |
| *DCAR_019805* | T | T | I | V | Y | Q | A | W | Y |
| *Aqcoe1G405000* | T | T | I | V | Y | Q | A | W | Y |
| *evm.TU.supercontig_19.60* | T | T | V | V | Y | Q | F | W | Y |
| *Vadar_g16772* | T | T | V | V | Y | Q | T | W | Y |
| *Prupe.2G263900* | T | T | V | V | Y | Q | A | W | Y |
| *MD07G1233400* | T | T | V | V | Y | - | - | W | Y |
| *MD01G1167300* | T | T | D | L | - | - | - | W | Y |
| *Fxa7Dg102232* | T | T | I | V | Y | Q | S | W | Y |
| *Fxa7Bg202436* | T | T | V | V | Y | Q | S | W | Y |
| *Fxa7Ag202526* | T | T | V | V | Y | Q | S | W | Y |
| *FvH4_7g25890* | T | T | V | V | Y | Q | S | W | Y |
| *Fxa7Cg102357* | T | T | V | V | Y | Q | S | W | Y |
| *29729.t000101* | T | T | V | V | Y | Q | A | W | Y |
| *Manes.07G140900* | T | T | V | V | Y | Q | A | W | Y |
| *CKAN_00912300* | T | T | V | V | Y | Q | A | W | Y |
| *LitulAlt.15G009500* | T | T | V | V | Y | Q | A | W | Y |
| *Anaoc.0016s1017* | T | T | I | V | Y | Q | A | W | Y |
| *Anaoc.0011s0947* | T | T | V | V | Y | Q | A | W | Y |
| *Ptrif.0004s0281* | T | T | I | V | Y | Q | A | W | Y |
| *Ciclev10032749m.g* | T | T | I | V | Y | Q | A | W | Y |
| *VIT_213s0067g02870* | T | T | I | V | Y | Q | A | W | Y |
| *Caden.06G174000* | T | T | I | V | Y | Q | S | W | Y |
| *NdRogue1H2.08G251000* | T | T | I | V | Y | Q | S | W | Y |
| *Qurub.08G299100* | T | T | I | V | Y | Q | S | W | Y |
| *CiPaw.02G198800* | T | T | I | V | Y | Q | A | W | Y |
| *Sapur.019G061200* | T | T | I | V | Y | Q | V | W | Y |
| *Podel.19G059600* | T | T | V | V | Y | Q | V | W | Y |
| *Potri.019G057800* | T | T | V | V | Y | Q | V | W | Y |
| *Thecc.10G181400* | T | T | V | V | Y | Q | A | W | Y |
| *Gohir.D04G012300* | T | T | V | V | Y | Q | A | W | Y |
| *Gohir.A05G403200* | T | T | V | V | Y | Q | A | W | Y |
| *Corav.Jeff.Hap2_g24033* | T | T | I | V | Y | Q | A | W | Y |
| *CamerWinkler.07G220100* | T | T | I | V | Y | Q | A | W | Y |
| *BPChr11G07344* | T | T | I | V | Y | Q | A | W | Y |
| *MyflaH2.07G002400* | T | T | I | V | Y | Q | A | W | Y |
| *Cocit.E0959* | T | T | I | V | Y | P | A | W | Y |
| *Eucgr.G03138* | T | T | I | V | Y | L | A | W | Y |
| *Lus10012878.g* | T | T | V | V | Y | Q | A | W | Y |
| *Lus10030529.g* | T | T | V | V | Y | Q | A | W | Y |
| *Sapof.03G134000* | T | T | V | V | Y | Q | A | W | Y |
| *FUN_018973* | T | T | V | V | Y | Q | A | W | Y |
| *Bevul.2G026900* | T | T | A | V | L | Q | S | W | Y |
| *EL10Ac2g02642* | T | T | A | V | L | Q | S | W | Y |
| *AH019282* | T | T | - | V | L | Q | S | W | Y |
| *Spov3_chr4.01990* | T | T | A | V | L | Q | S | W | Y |
| *AUR62024932* | T | T | A | V | L | Q | A | W | Y |
| *AUR62030612* | T | T | A | V | L | Q | A | W | Y |
| *Cucsa.143940* | T | T | V | V | Y | Q | A | W | Y |
| *ChfasH2.7G121000* | T | Y | I | V | Y | Q | A | W | Y |
| *CecanH2.4G235200* | T | Y | L | I | Y | Q | A | W | Y |
| *ChfasH2.7G204700* | T | Y | L | I | Y | Q | A | W | Y |
| *Lalb_Chr23g0272211* | - | F | L | V | Y | Q | A | W | Y |
| *Lalb_Chr08g0239161* | - | - | L | V | Y | Q | A | W | Y |
| *Medtr3g093980* | T | Y | L | V | Y | Q | A | W | Y |
| *Tp57577_TGAC_v2_gene16134* | T | Y | L | V | Y | Q | A | W | Y |
| *Vfaba.Hedin2.R1.2g141160* | T | Y | L | V | Y | Q | A | W | Y |
| *Ler.1DRT.3g060960* | T | Y | L | V | Y | Q | A | W | Y |
| *Lcu.2RBY.3g055360* | T | Y | L | V | Y | Q | A | W | Y |
| *Lj1g0023491* | N | Y | L | V | Y | Q | A | W | Y |
| *arahy.Tifrunner.gnm1.ann1.WYW97W* | K | Y | L | V | Y | Q | A | W | Y |

| | 36 | 48 | 95 | 97 | 106 | 109 | 113 | 190 | 191 |
|---|---|---|---|---|---|---|---|---|---|
| *arahy.Tifrunner.gnm1.ann1.SNS0I7* | K | Y | L | V | Y | Q | A | W | Y |
| *Vigun09g132600* | T | Y | L | V | Y | Q | A | W | Y |
| *Phcoc.09G145600* | T | Y | L | V | Y | Q | A | W | Y |
| *Pl09G0000157100.v1* | T | Y | L | V | Y | Q | A | W | Y |
| *PvUI111.09G144300* | T | Y | L | V | Y | Q | A | W | Y |
| *Phacu.WLD.009G079700* | T | Y | L | V | Y | Q | A | W | Y |
| *Glyma.04G222400* | T | Y | L | V | Y | Q | A | W | Y |
| *GlysoPI483463.04G180100* | T | Y | L | V | Y | Q | A | W | Y |
| *GlymaFiskIII.04G203400* | T | Y | L | V | Y | Q | A | W | Y |
| *GlymaLee.04G183300* | T | Y | L | V | Y | Q | A | W | Y |
| *Glyma.06G143000* | T | Y | L | V | Y | Q | A | W | Y |
| *GlysoPI483463.06G130600* | T | Y | L | V | Y | Q | A | W | Y |
| *GlymaFiskIII.06G136200* | T | Y | L | V | Y | Q | A | W | Y |
| *GlymaLee.06G131600* | T | Y | L | V | Y | Q | A | W | Y |

**Supplementary Figure 2**

```
MsCHI-II     1 MAA-SITAITVENLEYPAVVTSPVTGKSYFLGGAGERGLTIEGNFIKFTAIG  51
MtCHI-I      1 MALPSVTALEIENYAFPPTVKPPGSTNNFFLGGAGERGIQIQDKFVKFTAIG  52
AtCHIL       1 MGT-EM--VMVHEVPFPPQIIT---SKPLSLLGQGITDIEIHFLQVKFTAIG  46
OsCHIL       1 MGT-EIATVEVEGIPFPQEITV---SKPLSLLANGITDIEIHFLQIKYNAIG  48
GmCHIL       1 MAT-EE--VLVDEITYPTKITT---TKPLSLLGHGITDMEIHFIHVKFYSIG  46
SmCHIL       1 MEM-DP--------TFAQSIQSPSSSETLILLGHGITDMTIETIHVIFTKIG  43
MpCHIL       1 MAQ-GD--YAVDGIEFPATIQVPD-SPPLKFLGQGVTDMTIETILVKFTAIG  48

MsCHI-II    52 VYLEDIAVASLAAKWKGKSSEELLET-LDFYRDIISGPFEKLIRGSKIRELS 102
MtCHI-I     53 VYLQDIAVPYLAEKWKARSAHELTDT-VPFFRDIVTGPFEKFMRVTMILPLT 103
AtCHIL      47 VYLDPSDVKTHLDNWKGKTGKELAGD-DDFFDALASAEMEKVIRVVVIKEIK  97
OsCHIL      49 VYLEKDNVLAHLESWKGKKAEELVQD-DGFFQALVSAPVEKLLRIVVIKEIK  99
GmCHIL      47 VYLEPE-VVGHLDQFKGKSAKELEDN-EEFFNALISAPVEKFIRLVVIKEIK  96
SmCHIL      44 VYFAPQ-VKDHLQSFKCLPVSELLKDGSAFFQQLIQAPVSKLIKILLVKGQL  94
MpCHIL      49 VYASED-IKGHLKAWGGKASSDLTAEESGYFKDFVDAPVPKLLKIGVIKGIK  99

MsCHI-II   103 GPEYSRKVMENCVAHLKSVGTYGDAEAEAMQKFAEAFKPVNFPPGASVFYR- 153
MtCHI-I    104 GHQYSEKVSENCVAIWKSLGIYTDEEAKAIDKFVSVFKDETFPPGSSILFT- 154
AtCHIL      98 GAQYGVQLENTVRDRLAEEDKYEEEEETELEKVVGFFQSKYFKANSVITYHF 149
OsCHIL     100 GSQYGVQLESSVRDRLVSVDKYEEDEEEALEKVTEFFQSKYFKPNSVITFHF 151
GmCHIL      97 GAQYGVQIETAVRDRLAAEDKYEEEEEEALEKVIEFFQSKYFKKLSVITYHF 148
SmCHIL      95 GSQYASTIETSVRDRLAYDDKYEEDEEIALANLCEFFQSKKLEPNSTIVYSW 146
MpCHIL     100 GNQYGGTLESSIRDRLAYDDKYEEEEEAVVERIVEYLETKNLPPGSCMFIYW 151

MsCHI-II   154 QSPDGI--LGLSFSPDTSIPEKEAALIENKAVSSAVLETMI-GEHAVSPDLK 202
MtCHI-I    155 VSPKGLGSLTISFSKDGSIPEVETAVIENKLLSQAVLESMI-GAHGVSPAAK 205
AtCHIL     150 SAKDGI--CEIGFETEG--KEEEKLKVENANVVGMMQRWYLSGSRGVSPSTI 197
OsCHIL     152 PTTPGI--AEISFVTEG--KGEAKLTVENKNVAEMIQKWYLGGESAVSPTTV 199
GmCHIL     149 PANSAT--AEIVVSLEG--KEDSKYVIENANVVEAIKKWYLGGSSAVSSSTI 196
SmCHIL     147 PSSSSH--VEVFVHEEGS-KAPSSFIVNNENVSTSIIEWIL-GENSMTPSTV 194
MpCHIL     152 PTPDEI---KMSVSTDNTIPEKFDFSAVNKNVATYLLHWYM-GENAMSETTS 199

MsCHI-II   203 RCLAARLPALLNEGAFKIGN                                 222
MtCHI-I    206 QSLASRLSKLFKEGG-NANN                                 224
AtCHIL     198 VSIADSISAVLT--------                                 209
OsCHIL     200 KSLADQFAALLSA-------                                 212
GmCHIL     197 QSLASTFSQELSK-------                                 209
SmCHIL     195 ESVAKSIATEC---------                                 205
MpCHIL     200 AALAEGVVALLA--------                                 211
```

### Supplementary Figure 3

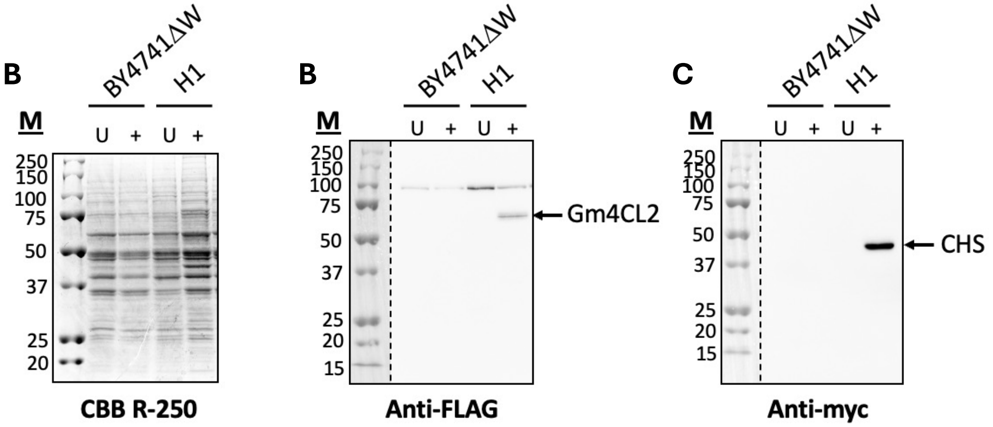

### Supplementary Figure 4

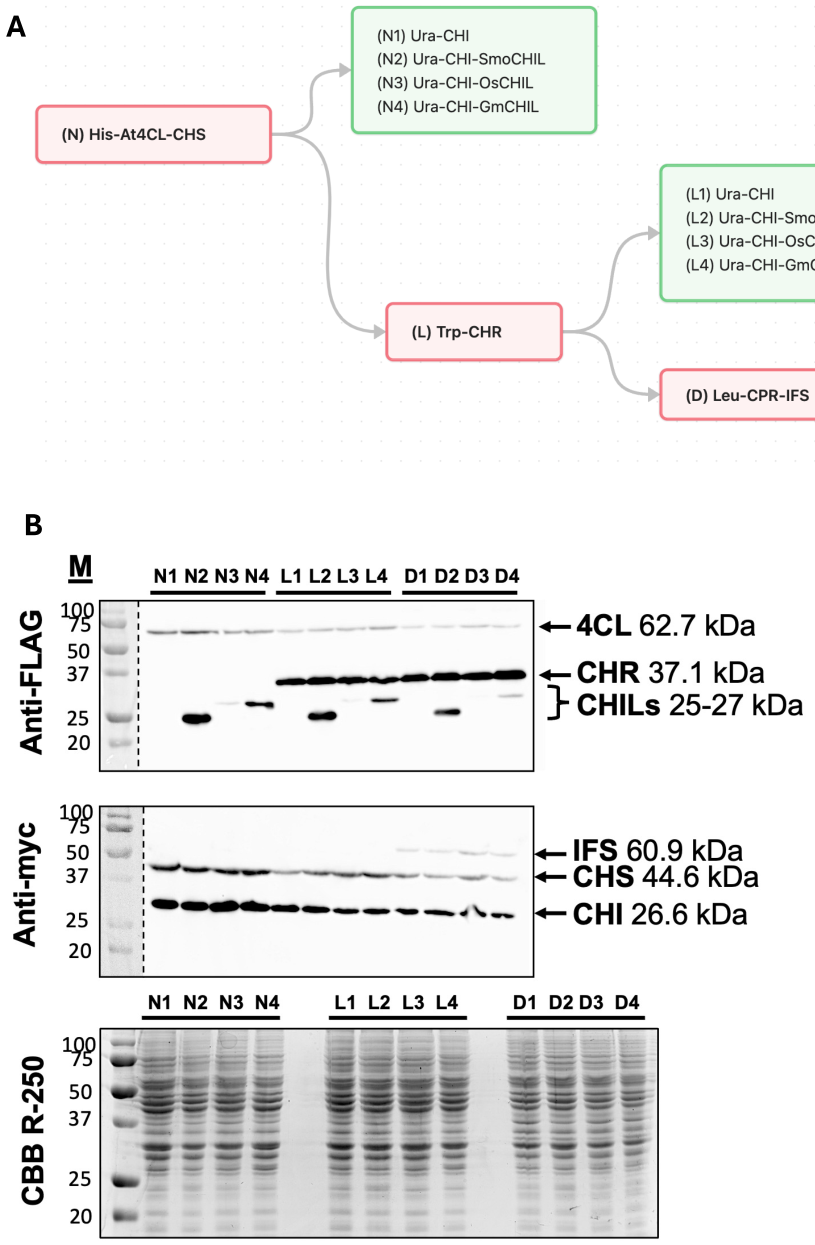

### Supplementary Figure 5

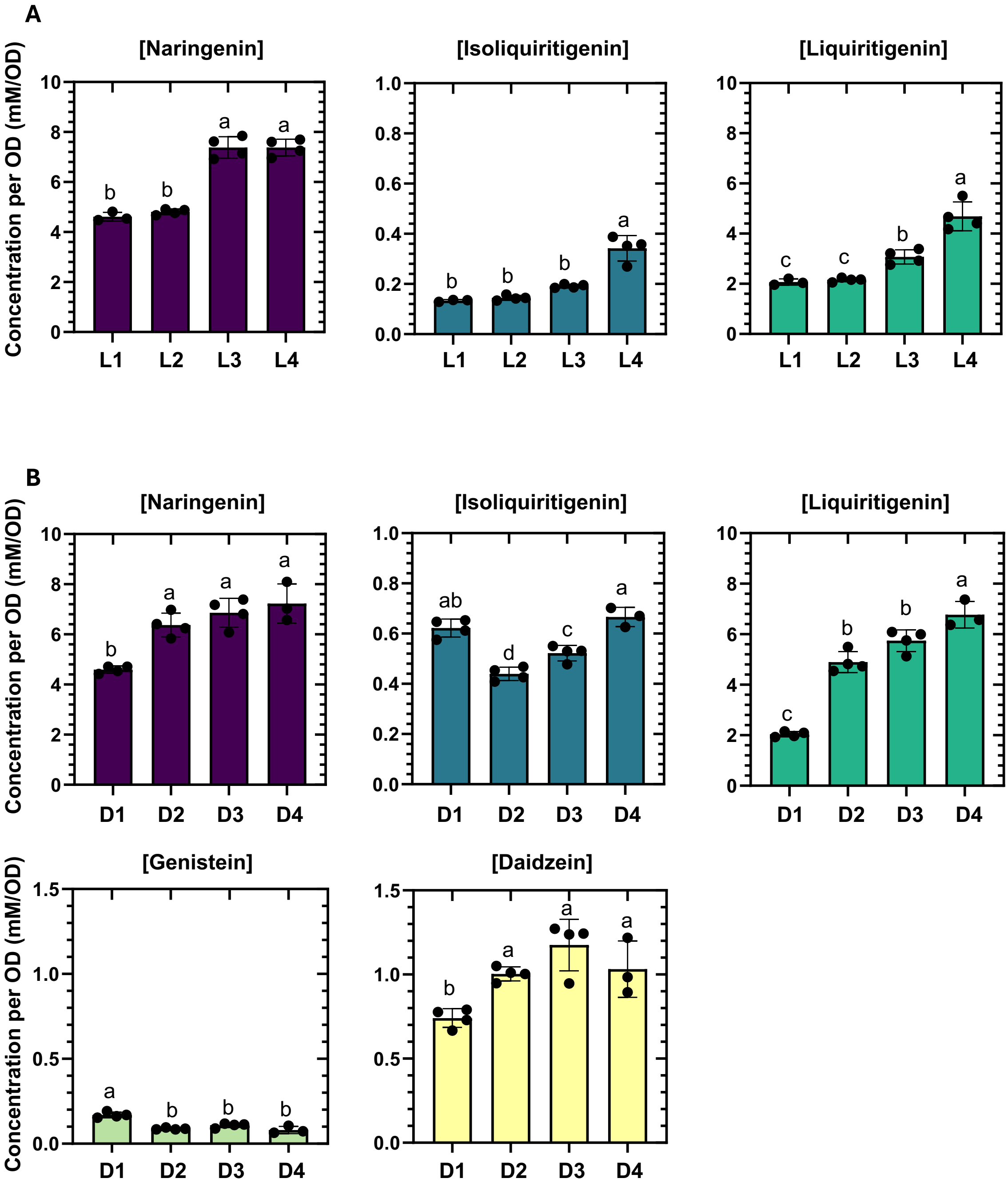
