## Supplementary Table for "Chalcone isomerase-like impedes the lactone shunt and enhances flux partitioning in a bifurcated pathway towards isoflavonoid biosynthesis"

**Supplementary Table 1.** List of yeast strains generated in this study in the BY4741ΔW background strain.

| Strain Name | Plasmids |
|---|---|
| H1 | pESC-*His-GAL1:CHS-myc-GAL10:Gm4CL2-FLAG* |
| H2 | pESC-*His-GAL1:CHS-myc-GAL10:At4CL1-FLAG* |
| N1 | pESC-*His-GAL1:CHS-myc-GAL10:At4CL1-FLAG*<br>pESC-*Ura-GAL1:CHI-myc* |
| N2 | pESC-*His-GAL1:CHS-myc-GAL10:At4CL1-FLAG*<br>pESC-*Ura-GAL1:CHI-myc-GAL10:SmCHIL-FLAG* |
| N3 | pESC-*His-GAL1:CHS-myc-GAL10:At4CL1-FLAG*<br>pESC-*Ura-GAL1:CHI-myc-GAL10:OsCHIL-FLAG* |
| N4 | pESC-*His-GAL1:CHS-myc-GAL10:At4CL1-FLAG*<br>pESC-*Ura-GAL1:CHI-myc-GAL10:GmCHIL-FLAG* |
| L0 | pESC-*His-GAL1:CHS-myc-GAL10:At4CL1-FLAG*<br>pESC-*Trp-GAL1:CHR-myc* |
| L1 | pESC-*His-GAL1:CHS-myc-GAL10:At4CL1-FLAG*<br>pESC-*Trp-GAL1:CHR-myc*<br>pESC-*Ura-GAL1:CHI-myc* |
| L2 | pESC-*His-GAL1:CHS-myc-GAL10:At4CL1-FLAG*<br>pESC-*Trp-GAL1:CHR-myc*<br>pESC-*Ura-GAL1:CHI-myc-GAL10:SmCHIL-FLAG* |
| L3 | pESC-*His-GAL1:CHS-myc-GAL10:At4CL1-FLAG*<br>pESC-*Trp-GAL1:CHR-myc*<br>pESC-*Ura-GAL1:CHI-myc-GAL10:OsCHIL-FLAG* |

| | |
|---|---|
| L4 | pESC-*His-GAL1:CHS-myc-GAL10:At4CL1-FLAG*<br>pESC-*Trp-GAL1:CHR-myc*<br>pESC-*Ura-GAL1:CHI-myc-GAL10:GmCHIL-FLAG* |
| D0 | pESC-*His-GAL1:CHS-myc-GAL10:At4CL1-FLAG*<br>pESC-*Trp-GAL1:CHR-myc*<br>pESC-*Leu2d-GAL1:IFS-myc-GAL10:CrCPR-FLAG* |
| D1 | pESC-*His-GAL1:CHS-myc-GAL10:At4CL1-FLAG*<br>pESC-*Trp-GAL1:CHR-myc*<br>pESC-*Leu2d-GAL1:IFS-myc-GAL10:CrCPR-FLAG*<br>pESC-*Ura-GAL1:CHI-myc* |
| D2 | pESC-*His-GAL1:CHS-myc-GAL10:At4CL1-FLAG*<br>pESC-*Trp-GAL1:CHR-myc*<br>pESC-*Leu2d-GAL1:IFS-myc-GAL10:CrCPR-FLAG*<br>pESC-*Ura-GAL1:CHI-myc-GAL10:SmCHIL-FLAG* |
| D3 | pESC-*His-GAL1:CHS-myc-GAL10:At4CL1-FLAG*<br>pESC-*Trp-GAL1:CHR-myc*<br>pESC-*Leu2d-GAL1:IFS-myc-GAL10:CrCPR-FLAG*<br>pESC-*Ura-GAL1:CHI-myc-GAL10:OsCHIL-FLAG* |
| D4 | pESC-*His-GAL1:CHS-myc-GAL10:At4CL1-FLAG*<br>pESC-*Trp-GAL1:CHR-myc*<br>pESC-*Leu2d-GAL1:IFS-myc-GAL10:CrCPR-FLAG*<br>pESC-*Ura-GAL1:CHI-myc-GAL10:GmCHIL-FLAG* |
